# Ultraviolet Dosage and Decontamination Efficacy was Widely Variable across 14 UV Devices after Testing a Dried Enveloped Ribonucleic Acid Virus Surrogate for SARS-CoV-2

**DOI:** 10.1101/2022.01.27.478063

**Authors:** Tony L. Buhr, Erica Borgers-Klonkowski, Bradford W. Gutting, Emlyn E. Hammer, Shelia M. Hamilton, Brett M. Huhman, Stuart L. Jackson, Neil L. Kennihan, Samuel D. Lilly, John D. Little, Brooke B. Luck, Emily A. Matuczinski, Charles T. Miller, Rachel E. Sides, Vanessa L. Yates, Alice A. Young

**Author notes:** Correspondence: Tony L. Buhr, PhD.

## Abstract

**Aims:** The dosages and efficacy of 14 ultraviolet (UV) decontamination technologies were measured against a SARS-CoV-2 surrogate virus that was dried on to different materials for lab and field testing.

**Methods and Results:** A live enveloped, ribonucleic acid (RNA) virus surrogate for SARS-CoV-2 was dried on stainless steel 304 (SS304), Navy Top Coat-painted SS304 (NTC), cardboard, polyurethane, polymethyl methacrylate (PMMA), and acrylonitrile butadiene styrene (ABS) at > 8.0 log_10_ plaque-forming units (PFU) per test coupon. The coupons were then exposed to UV radiation during both lab and field testing. Commercial and prototype UV-emitting devices were measured for efficacy: 4 handheld devices, 3 room/surface-disinfecting machines, 5 air-disinfection devices, and 2 larger custom-made machines. UV device dosages ranged from 0.01-729 mJ cm^-2^. Anti-viral efficacy among the different UV devices ranged from no decontamination up to nearly achieving sterilization. Importantly, cardboard required far more dosage than SS304.

**Conclusions:** Enormous variability in dosage and efficacy was measured among the different UV devices. Porous materials limit the utility of UV decontamination.

**Significance and Impact of the Study:** UV devices have wide variability in dosages, efficacy, hazards, and UV output over time indicating that each UV device needs independent technical measurement and assessment for product development, prior to, and during use.

## INTRODUCTION

Tremendous attention was directed at the subject of UV decontamination during the COVID-19 pandemic, although UV devices are best used to augment other sanitation techniques rather than for stand-alone decontamination as successful use of UV depends on various environmental and technical factors (Raiszadeh and Adeli 2020; Memarzadeh 2021). Numerous devices that incorporate UV sources including handheld, room decontamination devices, and water treatment devices are available on the market to decontaminate air, water, and surface materials. Variability in UV devices is extensive and includes differences in electronics, bulbs, power, and product designs. The distance from UV sources at which decontamination/inactivation occurs is also widely variable ranging from a couple of centimeters to a couple of meters. UV sources also differ and include mercury (Hg), krypton chloride (KrCl), xenon (Xe), and various light emitting diodes (LED), which range in wavelength, and there are several different manufacturers. Additionally, although Hg bulbs are the most common, Hg bulb dosage significantly varies over time after the Hg bulb is turned on. Hg also comes with the risk of toxicity, although that risk is similar to fluorescent light bulbs. The variability in these decontamination devices is further complicated by variability in test methods, which include different virus preparation methods, tests with unpurified vs. purified virus, tests with wet virus or dried virus, presence of organic debris, and differences in porosity of surface materials.

Assessments of UV for decontamination must also take into account maintenance since UV sources need to be cleaned in order to maintain dosage (United States Food and Drug Administration 2021; United States Environmental Protection Agency 2021).

UV radiation, particularly UV-C, is a known microbe disinfectant for air, water, and nonporous surfaces (United States Food and Drug Administration 2021; United States Environmental Protection Agency 2021). UV-C primarily inactivates microbes including viruses if they are directly exposed to the UV radiation. Therefore inactivation is far less effective if a microbe is associated with soil, dust, oils, any type of host cell debris, or if it is embedded in porous materials (United States Food and Drug Administration 2021). This is particularly relevant for obligate pathogens like viruses which are naturally associated with host cell components and body fluids: mucus in the case of respiratory virus like SARS-CoV-2 (Aps and Marten 2005; Heimbuch et al. 2013; Vejerano and Marr 2018). The effectiveness of UV-C lamps in inactivating environmentally relevant SARS-CoV-2 virus is unknown because there is limited consistent and/or reliable published data about the wavelength, dose, and duration of UV-C radiation required to inactivate the SARS-CoV-2 virus, particularly in its natural (unpurified) state (United States Food and Drug Administration 2021; United States Environmental Protection Agency 2021). This is true of all viruses because UV efficacy is further complicated by the fact that methods for virus preparation and testing, particularly enveloped viruses, are highly variable among laboratories (Hadi et al. 2020). Purified enveloped viruses are often tested in laboratories, even though these viruses only exist naturally when associated with host cell components and debris in nature, and they can be compromised during purification (Cox and Wathes 1995).

The stability of viral particles in the environment depends on temperature and humidity, as well as characteristics of the virus itself as it is shed from the host (Cox and Wathes 1995). Human respiratory droplets are mainly composed of mucus (salt, mucin glycoprotein, and lipids (surfactants)), and these components can shield the virus from UV and affect decontamination kinetics (Vejerano and Marr 2018; Hadi et al. 2020). The mean size of infectious, respirable particles is 4 um for a 50% probability of a thoracic deep lung deposition (Brown et al. 2013).

Particles >10 um don’t go past the larynx and particles <1 um have lower probabilities of deposition (Brown et al. 2013; Hofer et al. 2021). Measurements of SARS-CoV-2 respiratory droplets are typically 0-1 virions per speech particle, and the water in SARS-CoV-2 respiratory particles evaporates within seconds to generate dry particles around 4 um, right at the respirable size range (Stadnytskyi et al. 2020). Hence, the virus respiratory size is much larger than the size of a naked coronavirus, which are 78 nm for SARS-CoV (Goldsmith et al. 2004) and can range from 50-200 nm (Masters 2006). A volume/volume calculation with a 78 nm virus and 4 um particle equates to >99.999% mucus and <0.001% virus per particle and sets a minimum target on the ratio of debris to virus expected for decontamination testing. This only accounts for debris in respiratory particles and does not account for additional debris that might be found on surfaces. In addition, enveloped viruses are more stable in dry conditions compared to wet environments (Cox and Wathes 1995; Chan et al. 2011; Buhr et al. 2020; Hadi et al. 2020; van Doremalen et al., 2013), and drying viruses via lyophilization is frequently used to stabilize virus for long-term storage (Greiff et al. 1954; Greiff and Richtel 1966; Malenovska 2014). Hence decontamination kinetics can also be greatly influenced depending on whether the test microbes are wet or dry. Rhinotillexis (nose-picking) creates additional environmental loads of infectious virus, which is also composed of mucus mixed with unpurified virus and varying levels of free water (Hendley et al. 1973; Weber et al. 2008).

In addition to methods gaps to define, characterize, and standardize SARS-CoV-2 virus debris composition and drying, standardized methods for reproducibly preparing high titers (>10 log_10_ of virus ml^-1^ of culture medium at the time of virus harvest) of SARS-CoV-2 for testing without artificial post-harvest cleaning and concentration steps are needed for statistical confidence and to match virus levels in the environment. Furthermore, there were/are urgent needs during the COVID-19 pandemic to test decontamination devices, like UV, in field tests outside of laboratory containment. Viruses that fall under higher World Health Organization (WHO) biosafety level (BSL) classifications such as SARS-CoV-2 (BSL-3) and its BSL-2 surrogate coronaviruses (ASTM International 2020) cannot be widely used in field tests because of cost, time, and safety constraints. For field testing, the enveloped virus surrogate Ф6 was previously used to make “live/dead” Ф6 test indicators to directly test and compare decontamination efficacy across lab and field tests (Buhr et al. 2020).

Surrogates are often used as models in studies of decontamination for highly infectious pathogens. Bacteriophages are useful for this purpose as they are similar in terms of morphology, behavior in the environment, and surface properties, but are BSL-1 and easier to isolate at high titers (>10 log_10_ ml^-1^ without virus purification and concentration) for testing than mammalian viruses (Gallandat and Lantagne 2017). Among bacteriophages, Ф6 has been identified as a preferred surrogate for enveloped viruses including influenza and SARS-CoV-2 (Bibby et al. 2015; Gallandat and Lantagne 2017; Vejerano and Marr 2018; Buhr et al. 2020; Fedorenko et al. 2020). *Pseudomonas* virus Ф6 is a BSL-1 enveloped RNA virus originally isolated in a bean field as a lytic virus that infects the plant pathogenic bacterium *Pseudomonas syringae* pathovar *phaseolicola* (Vidaver et al. 1973; Van Etten et al. 1976; Mindich 2004). The Ф6 envelope structure is similar to many other enveloped viruses as the envelope consists of a glycoprotein/protein-embedded lipid membrane, and the host cell has similar temperature sensitivity to mammalian cells at around 40°C. This is important since the envelope components are considered a target for inactivation by many different decontaminants including UV radiation, particularly at 222 nm (McDonnell and Burke 2011; Wiggington et al. 2012; Hadi et al. 2020; United States Food and Drug Administration 2021). Φ6 is a 13.5 kb double-stranded RNA (dsRNA) phage (Mindich 2004), and spherical (80-100 nm diameter) with structural similarity to coronaviruses (50-200 nm diameter). The 13.5 kb dsRNA genome, the equivalent of 27 kb of single-stranded RNA (ssRNA), is comparable to the 26-32 kb of ssRNA in coronaviruses. In theory, a surrogate virus should have a similar number of adjacent pyrimidines compared to SARS-CoV-2 since pyrimidine dimerization is considered an important mechanism of UV inactivation (e.g. Heßling et al. 2020). Based on pyrimidine target numbers only, Ф6 (6,613 adjacent pyrimidine pairs) and SARS-CoV-2 (7,600 pairs) should have similar UV sensitivity, although ssRNA may be slightly more sensitive than dsRNA due to the potential for repair of dsRNA by the undamaged strand (Tseng and Li 2005). Hence sequence data alone theoretically implies that Ф6 inactivation goals should be similar to or slightly more conservative than SARS-CoV-2. Separately, it is currently difficult to compare UV efficacy both within and across different viruses based on existing data because experimental tests are highly variable across different labs and studies (Hadi et al. 2020). Overall, the sequence comparison between the two viruses is likely moot because debris, drying, and porosity of respiratory particles and/or contaminated surfaces have dominant impacts on decontamination kinetics particularly when the amount of debris is >99.999% relative to virus (United States Food and Drug Administration 2021; United States Environmental Protection Agency 2021). Furthermore, practical confidence that test methods approach the challenge of field conditions is needed from field decontamination testing in order to increase confidence in devices to be employed by end users (Hamilton et al. 2013; Buhr et al. 2015, 2016).

Here Ф6 was prepared at 11.0 ±0.2 log_10_ PFU ml^-1^ without post-harvest processing or concentration steps, and then dried on to different materials for >24 hours (h) to make BSL-1 live/dead enveloped virus test indicators at ≥ 8.0 log_10_ PFU coupon^-1^. Numerous UV devices were tested in both lab and field trials for both screening and iterative UV product improvement. As a BSL-1 surrogate, Ф6 is useful for the generation of baseline decontamination data for enveloped viruses, particularly during the COVID-19 pandemic when results from field decontamination methods and procedures were urgently needed. It is recognized that the limitation of Φ6 testing is that a correlation test with ≥5 independent batches of unpurified mammalian coronavirus at ≥ 8.0 log_10_ PFU coupon^-1^ is needed. The current limitation of such a correlation test is linked to limitations with coronavirus preparation and test methods. COVID- 19 mucus is significantly thicker than healthy mucus (Kratochvil et al. 2022), but not yet defined for virus inactivation testing. This natural microbial protectant will need standardized and combined with virus to generate practical confidence for SARS-CoV-2 inactivation data. In addition, titers of lab-prepared coronavirus needs to be significantly higher in order to meet the virus load that has been measured and is expected in the environment. While there are limitations in this study that center around the need for correlation testing with multiple, independent batches of unpurified, >99.999%-debris (mucus) laden, dried coronavirus at ≥ 8.0 log_10_ per test, the Ф6 test met the quantitative, practical objectives, and was the most conservative live/dead enveloped virus test known for field testing during COVID-19 and for screening/selecting decontamination equipment and technologies (Buhr et al. 2020).

## MATERIALS AND METHODS

### Φ6 and Host Cell Preparations

Virus and host cell preparation was previously described (Buhr et al. 2020). Φ6 and its host organism *P. syringae* pathovar *phaseolicola* HB10Y (HB10Y), causal agent of halo blight of the common bean, *Phaseolus vulgaris*, were isolated in Spain. Both were a kind gift from Dr. Leonard Mindich at Rutgers University, New Jersey Medical School. HB10Y was prepared by inoculating 100-200 ml of 3% tryptic soy broth (TSB; Fluka PN#T8907-1KG) in a 1-liter (l) smooth-bottom Erlenmeyer flask with a high efficiency particulate air (HEPA) filter cap.

Cultures were incubated at 26±2°C, 200 revolutions (rev) minute (min)^-1^ for 20±2 h. 11.1 ml of 100% glycerol (Sigma PN #G7757-500ML) was added per 100 ml of host culture. Final concentration of glycerol was 10%. One-ml aliquots of HB10Y were pipetted into screw-cap microfuge tubes with O-rings and stored at -80°C. HB10Y samples were titered prior to freezing by serially diluting samples in 10 millimolar (mM) of 4-(2-hydroxyethyl)-1- piperazineethanesulfonic acid (HEPES, Sigma PN#H4034-100G) + 10% Sucrose (Sigma PN #S7903-250G), pH 7.0, and plating on tryptic soy agar (TSA; Hardy Diagnostics, Santa Maria, CA). Plates were inverted and incubated at 26±2°C for 48±2 h to show titers of ∼10^9^ cells ml^-1^. After freezing, tubes were thawed at room temperature (RT, 22±3°C), serially diluted, and plated to show sustained viability after long-term storage at -80°C.

Ф6 was prepared after inoculating broth cultures of HB10Y. A frozen stock prep of HB10Y was thawed at 22±3°C. HB10Y was added either directly from a frozen stock or by transferring a single colony from a streaked TSA plate to 200 ml of 3% TSB in a 1-l smooth-bottom Erlenmeyer flask with a HEPA cap and incubated at 26±2°C, 200 rev min^-1^ overnight. Cells were then diluted and grown to mid-log-phase (3-5e8 cells ml^-1^). The host flask was inoculated with 0.5-1 ml of Φ6 at a stock concentration of 11±0.2 log_10_ PFU ml^-1^. The culture was incubated at 26±2°C, 200 rev min^-1^ for 24±2 h. The Ф6 preparation was stored at 4°C until after titering was completed. After titer determination was completed, then 1-1.3 ml volumes were aliquoted into 1.5-ml screw-cap tubes with O-rings, inverted, and stored at -80°C.

### Coupon Materials and Sterilization

2 centimeter (cm) x 2 cm coupons of different test materials were inoculated with ≥8.0 log_10_ PFU Φ6 virus inoculum (Buhr et al. 2020). Materials for inoculation included stainless steel 304 (SS304) (20 gauge with a 2B finish from Cardinal Scientific), SS304 coupons painted with Navy Top Coat (NTC) (26 gauge SS304 with a 2B finish primed with N-6237 and top coated with MIL-PRF-24635B, 4-6 mils from Coatings Group at the University of Dayton Research Institute (Dayton, OH, USA)), acrylonitrile butadiene styrene (ABS) plastic (flat black coupon from Cardinal Scientific), polymethyl methacrylate (PMMA) plastic (keyboard keys from Hewlett-Packard computer keyboards), polyurethane (clear polyurethane national stock number 9330-01-541-8524X3), and cardboard (hand-cut from a Corrugated Recycles, new, single wall, cardboard box (Davis Core & PAD, Cave Spring, Ga, USA) with thickness of 0.16 inch). ABS and PMMA plastics are often used for computer keyboards. The plastics and SS304 represent non-porous materials. NTC represent semi-porous surfaces found on military ships.

Cardboard represents porous materials used in shipping although it is not as porous as fabrics or carpeting.

For sterilization, SS304 and NTC coupons were rinsed with 18 mega-Ohm-cm, de- ionized water, placed on absorbent paper in an autoclave-safe container, and autoclaved for 30 min at 121°C, 100 kilopascals. PMMA keyboard keys were removed, trimmed, cleaned with soap, then rinsed with de-ionized water, and wrapped in aluminum foil. ABS coupons were similarly rinsed with de-ionized water and wrapped in foil. Cardboard coupons were devoid of noticeable debris, flaws, and ink, and were wrapped in foil. After wrapping in foil, the PMMA keyboard keys, ABS, and cardboard were all sterilized via hot, humid air at 95°C and 90% relative humidity (RH) for 4 h. Polyurethane coupons, having been pre-cut, were soaked in ethanol to remove ink residue left over from the cutting process. They were then rinsed with de- ionized water, sterilized via immersion in 70% ethanol for greater than 20 min, and allowed to dry. All sterilized coupons were stored in sterile containers until used.

### Coupon Inoculation, Extraction, and Quantitation

Five independent preparations of Φ6 were removed from -80°C storage and thawed at 22±3°C. Working inoculum was prepared by transferring stock Φ6 into 50-ml conical tubes containing 10mM HEPES + 10% Sucrose pH 7.0 with a final concentration of ∼9 log_10_ PFU ml^-1^. Coupons were inoculated with 0.1 ml of Φ6 working inoculum and subsequently held at 22±3°C for greater than 24 h to dry and adhere to the material. The PMMA keyboard keys were slightly slanted. Therefore, during inoculation and drying, the keys were positioned in a sterilized surface which was elevated on an incline via slats to provide a level inoculation surface.

Once the inoculum had dried onto the coupons, they were exposed to UV from the candidate devices. Specific parameters for testing the individual devices varied but coupon number and preparation prior to testing was maintained across all experiments. For each test, five individual coupons were included for each of the test materials (SS304, cardboard, NTC, polyurethane, and either PMMA plastic keyboard keys or ABS plastic). Each coupon was inoculated with one of five independent virus preparations as described above. Extraction and shipping control coupons (inoculated and transported to the testing sites but not exposed to UV radiation) as well as negative control coupons not inoculated with virus were also included for every experiment. Finally, the Φ6 virus inoculum used to prepare the coupons was maintained at RT from the date of coupon inoculation through the test, and viral titer was measured at the conclusion of test exposures for each experiment.

To increase confidence in decontamination results and to conservatively estimate decontamination requirements for enveloped virus in its native state, enveloped virus test coupons were prepared to be protected similar to a natural virus without interfering with the virus assay. Thus, Φ6 virus was unpurified to maintain natural stabilization with host cell debris and was diluted in a 10% sucrose solution to mimic the presence of carbohydrates in mucus without inhibiting the decontamination assay (Brakke 1951; Malenovska 2014; Buhr et al. 2020; Hadi et al. 2020; Stadnytskyi et al. 2020). In addition, enveloped virus was dried on coupons prior to testing since SARS-CoV-2 respiratory particles evaporate within seconds to generate dry particles, and drying on fomites is also historically documented as a route of infection for enveloped virus (Fenn 2001; Malenovska 2014; Hadi et al. 2020; Stadnytskyi et al. 2020).

After UV exposure, virus was extracted from both test and control coupons (**Figure 1**) and plated in <25 min using a Φ6 extraction and overlay procedure that was previously described (Buhr et al. 2020). For Φ6 extraction from materials (coupons), 5 ml of 10mM HEPES + 10% sucrose pH7 were added to each conical tube with a virus-inoculated coupon and vortexed for 2 min. After vortexing, 5 ml of HB10Y log-phase culture (confirmed with real-time Coulter Multisizer analysis) were added and allowed to infect at RT for 15 min, followed by 2 min of vortexing. Each sample was serially diluted, from -2 to -6, in 900 µl of 10 mM HEPES + 10% sucrose pH7. For each Φ6 dilution, from -1 to -6, 200 µl were transferred into individual tubes containing 200 µl log-phase HB10Y. Then 200 µl of those Φ6/HB10Y mixtures were added to individual TSB overlay tubes, poured onto individual TSA plates, and allowed to solidify for ≥30 min. Additionally, 1,000 µl was transferred from the 50 ml sample conical tube directly to a TSB overlay tube, and the remaining 8.3 ml was poured onto two TSA plates, and also allowed to solidify for ≥30 min. Solidified plates were then inverted, incubated for 20+/-2 h at 26°C, and quantified. Plates were incubated an additional 24 h, RT and quantified a final time.

**Figure 1.**
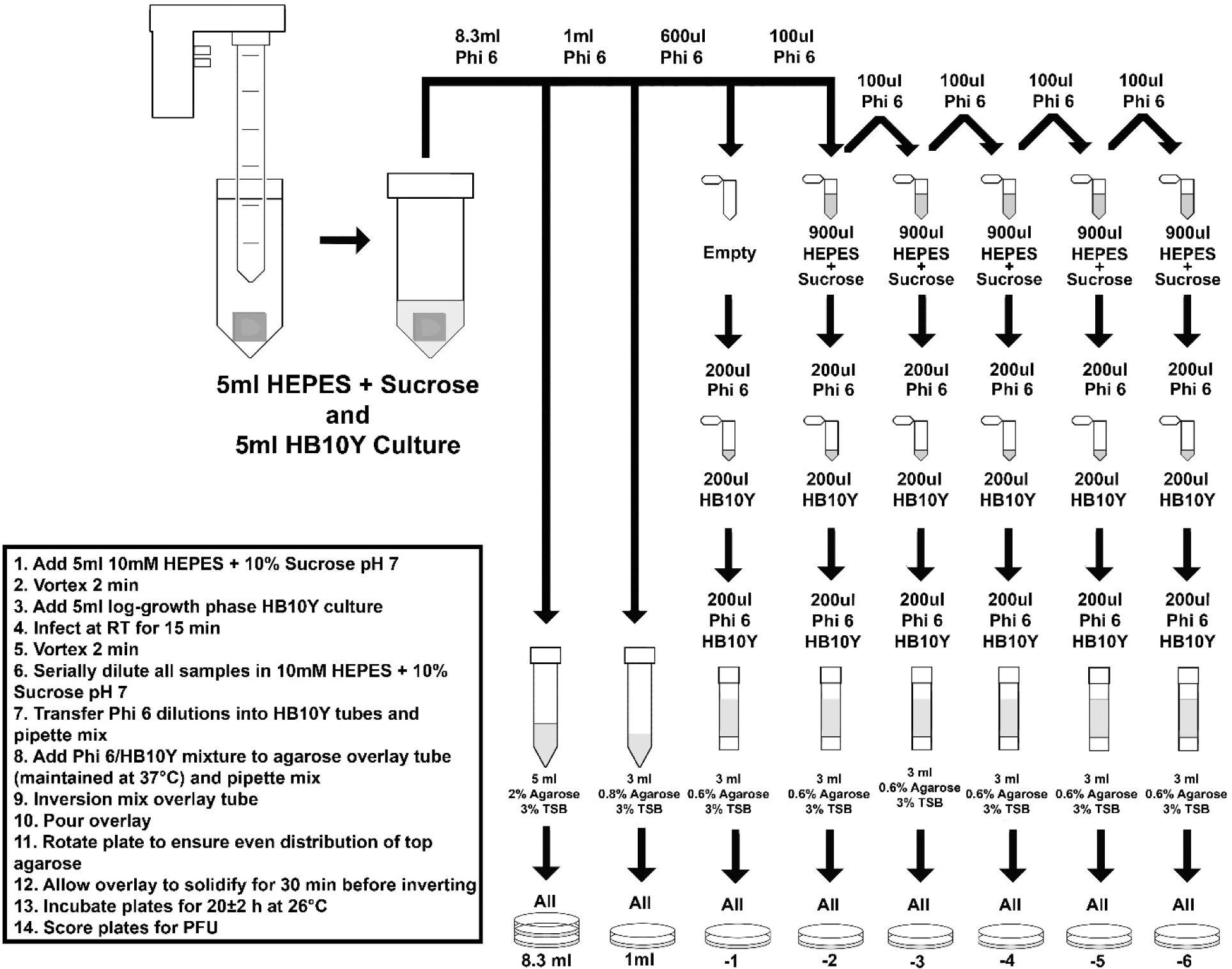
Φ6 extraction procedure for test samples and non-treated controls.

Quantitation and calculations of survival were performed as previously described (Buhr et al. 2020). An important difference between virus and prior spore quantitation is that virus and spore inoculum dried on to coupons was stable. However, titers of virus controls stored in solution were unstable and highly variable whereas spore controls were stable in solution.

Therefore, virus inoculation titers were defined as 100% extraction, or maximum recoverable virus, and used to calculate extraction efficiency for each material.

Virus survival and log_10_ reduction were then quantified using ASTM standard practices E3092 and E3178 except plaque forming units (PFU) replaced colony forming units (CFU). The sample mean and sample standard deviations were calculated for each set of samples. The virus inoculation titer adjusted for the volume and dilution difference served as the 100% recovery reference value for calculating the virus survival after decontamination. The extraction percentage was an arithmetic calculation not a log_10_ calculation. The extraction percentage was calculated by dividing the total number of infectious virus (PFU) extracted from the virus-inoculated control coupons for each material type by the total number of inoculated virus (PFU) (divided by 100 to adjust for 0.1 ml of viral inoculum deposited on each coupon and the 10 ml extraction volume). An average extraction efficiency was calculated and recorded for each type of test material for each test day. The number of surviving PFU for each test coupon was corrected for extraction efficiency by dividing the number of surviving PFU by the extraction percentage to determine the total number of PFU ml^-1^. The virus survival of each test coupon (corrected for extraction efficiency) in PFU ml^-1^ was multiplied by 10 to account for the 10 ml total volume in each tube with extraction buffer. This gave the total number of surviving PFU per sample. The virus survival of each control coupon in PFU ml^-1^ was also multiplied by 10 to account for the 10 ml of extraction volume. Each number was converted to 1og_10_. Since the log_10_ of 0 is infinite and for simplicity, the number ‘1’ was added to each number prior to converting to log_10_. The 1og_10_ mean and log_10_ standard deviation was calculated for infectious virus for each coupon material. The 1og_10_ reduction mean and standard deviation was calculated for each coupon material. For the 1og_10_ reduction, the 1og_10_ mean of each coupon material was subtracted from the 1og_10_ mean of the virus titer (minus 1 to adjust for 0.1 ml of viral inoculum deposited on to each coupon), the 100% recovery reference value. For the 1og_10_ reduction standard deviation, the square root of the result of the 1og_10_ survival standard deviation was taken for each coupon squared; divided by the number of independent samples plus the 1og_10_ survival standard deviation of the virus inoculum titer squared; divided by the number of independent samples.

### Spectroscopic Analysis Hardware and Calibration

The primary spectrometer used for this work was the Ocean Optics Maya 2000 Pro, which measured optical spectra from 180 – 630 nm with an average bin size of 0.22 nm across the measurable spectrum. The distribution is not strictly linear, but can be specifically determined as necessary for data processing. The spectrometer was used with a fiber bundle (BFL200HS02), which incorporates seven Φ200-μm core fibers into a single high-OH package. This enables the measurement of sources with low output so the spectrometer can both retain a high signal-to-noise ratio and enable the use of a cosine corrector (CCSA2) for most measurements.

The Maya 2000 Pro spectrometer was calibrated using a Cathodeon R48 Deuterium Lamp, serial number CH5627. The spectral irradiance from this lamp is in units of mW•m^-2^•nm^-1^ in 5 nm intervals from 200–400 nm. To perform the calibration, the lamp is mounted vertically and positioned so that a horizontal line through the center of the area to be irradiated passes through the center of the lamp emission area, as well as perpendicular to the lamp window. The calibration refers to the spectral irradiance over an approximately 10 mm^2^ area in a vertical plane located at a distance of 200 mm from the outside surface of the output window on the lamp. The lamp is operated from a 300-mA power supply and must be operated continuously for 30 min prior to recording data on the spectrometer.

The spectrometer was mounted on an optical table, with a three-axis linear translation stage (Thorlabs LTS300) used to enable precision alignment between the spectrometer fiber sensor head and the source of interest. The three-axis system is capable of measuring a 300 mm x 300 mm x 300 mm volume with computer automation using a process-controlled script via the Thorlabs Kinesis software. The data acquisition software used National Instruments LabVIEW for all aspects except direct control of the translation stages. All of the data was written to a single Technical Data Management Streaming data file for post-processing, which enabled all of the measurements to have a common time base for analysis. Post-processing was accomplished with the Jupyter software environment with discrete Python code blocks to allow for processing of specific sources as needed. The raw TDMS data file is loaded into a cache file on the processing server, and a series of factors and calibrations are applied to prepare the raw data for analysis. Static measurements are relatively simple, as the position is fixed and no further analysis is required. Sweeps in a two-dimensional space with the translation stages requires synchronization of the position with digital fiducial markers to construct an image of the measured plane at a given distance from the source.

## RESULTS

A focus of this work was to generate information for screening field devices and to provide feedback for iterative product improvement. Some data for prototypes were deliberately omitted since all prototypes were in the process of iterative improvement. The variability of UV emission from different test bulbs and devices (**Figure 2)** was enough by itself to justify the need for high confidence, quantitative, field test standards to guide product screening and iterative improvements in UV devices. Differences in intensity, dosages and virucidal efficacy among devices were additional justifications for standardized field testing described later.

**Figure 2.**
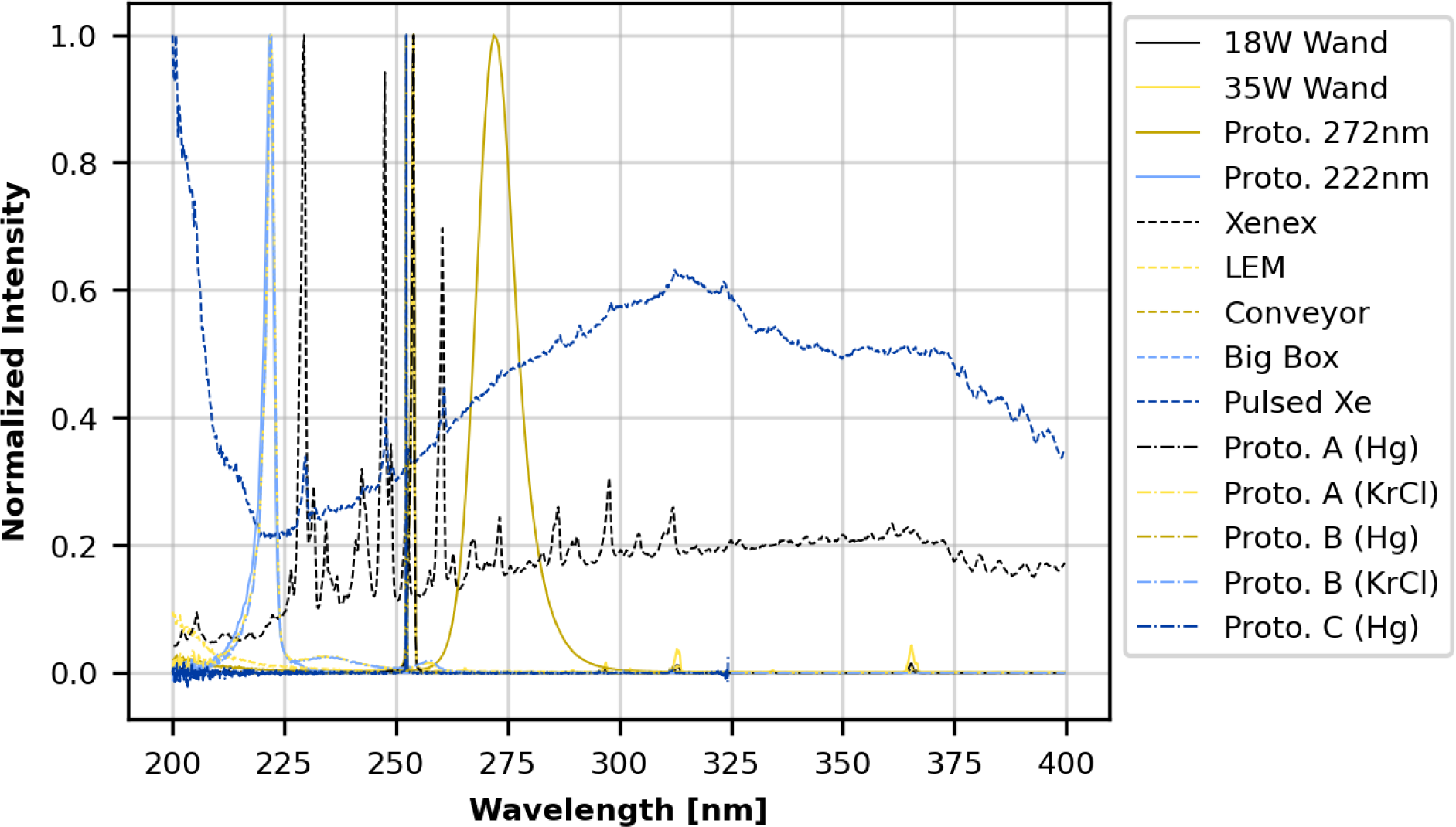
Normalized wavelength emission spectra for four handheld devices (18 watt wand (solid black line); 35 watt wand (solid light gold); 272 nm prototype (solid dark gold); 222 nm prototype (solid light blue)), five room devices (Xenex (dashed black); LEM (dashed light gold); medium conveyer (dashed dark gold); big box (dashed light blue); pulsed Xe (dashed dark blue), and five prototype devices (prototype A - Hg (dot-dashed black)); prototype A - KrCl (dot- dashed light gold); prototype B - Hg (dot-dashed dark gold); prototype B - KrCl (dot-dashed light blue); prototype C (dot-dashed dark blue)).

Extraction of viable virus from each non-treated control sample demonstrated that extraction efficiency was consistent across materials for 4-14 days (d) after coupon inoculation with >8 log_10_ virus and subsequent drying (**Table 1**). This was an important goal to meet in order to demonstrate that the method could generate reproducible results for tests at multiple field test sites outside of laboratory containment. The selection of materials to be inoculated for different devices was dependent on the intended use of different devices, the iteration of testing, and the availability of different materials, noting that different materials were available at different times during testing because of procurement limitations during COVID-19. After initial virus testing with keyboard keys, flat ABS coupons were procured. The keyboard keys were later identified as PMMA plastic rather than ABS plastic through Fourier Transform Infrared Spectroscopy, although both types of plastics are used for keyboards and both are non- porous, hard plastics.

**Table 1.**
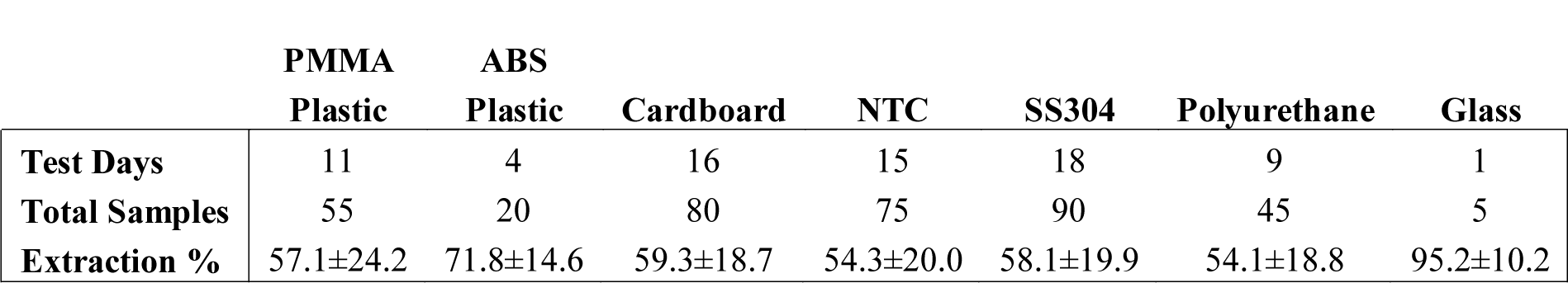
Average extraction efficiency of Φ6 virus 4-14 d after drying on different materials.

|  | PMMA<br>Plastic | ABS<br>Plastic | Cardboard | NTC | SS304 | Polyurethane | Glass |
| --- | --- | --- | --- | --- | --- | --- | --- |
| Test Days | 11 | 4 | 16 | 15 | 18 | 9 | 1 |
| Total Samples | 55 | 20 | 80 | 75 | 90 | 45 | 5 |
| Extraction % | 57.1±24.2 | 71.8±14.6 | 59.3±18.7 | 54.3±20.0 | 58.1±19.9 | 54.1±18.8 | 95.2±10.2 |

### Commercial handheld devices (18-watt, 35-watt)

Two commercial handheld devices were acquired and tested, each within a custom test apparatus. The first was the GermAwayUV 18W Handheld UV-C Surface Sanitizer (SKU 202110, bulb SKU 195317, CureUV, Delray Beach, FL, USA), a 120V/60Hz device containing two 12.7-cm long, U-shaped (Hg) UV bulbs emitting 254 nm UV-C radiaton (**Figure 3A**). An average intensity of 7.61 mW cm^-2^ was measured within a decontamination footprint of 4.47 cm x 5.39 cm at a 5-cm standoff distance from the bulb (heat map of UV coverage is shown in **Figure 3C)**. The second device was the GermAway UV Premier 35W Handheld UV-C Surface Sanitizer (PN14-110-800-100, EPA Product No. 94850-DV-6, CureUV, Delray Beach, FL, USA), 120V/60Hz handheld containing two Hg bulbs that emit 254 nm UV-C radiation, with reflective material positioned within the unit to enhance UV coverage (**Figure 3B**). The twin tube bulbs spanned a length of 22.5 cm. The 35W device provided an average intensity of 6.95 mW cm^-2^ at 5-cm standoff distance from the bulb (**Figure 3D**). The 35W handheld was later discovered to contain ineffective ballasts (P/N 14-110-800-100), which negatively impacted results.

**Figure 3.**
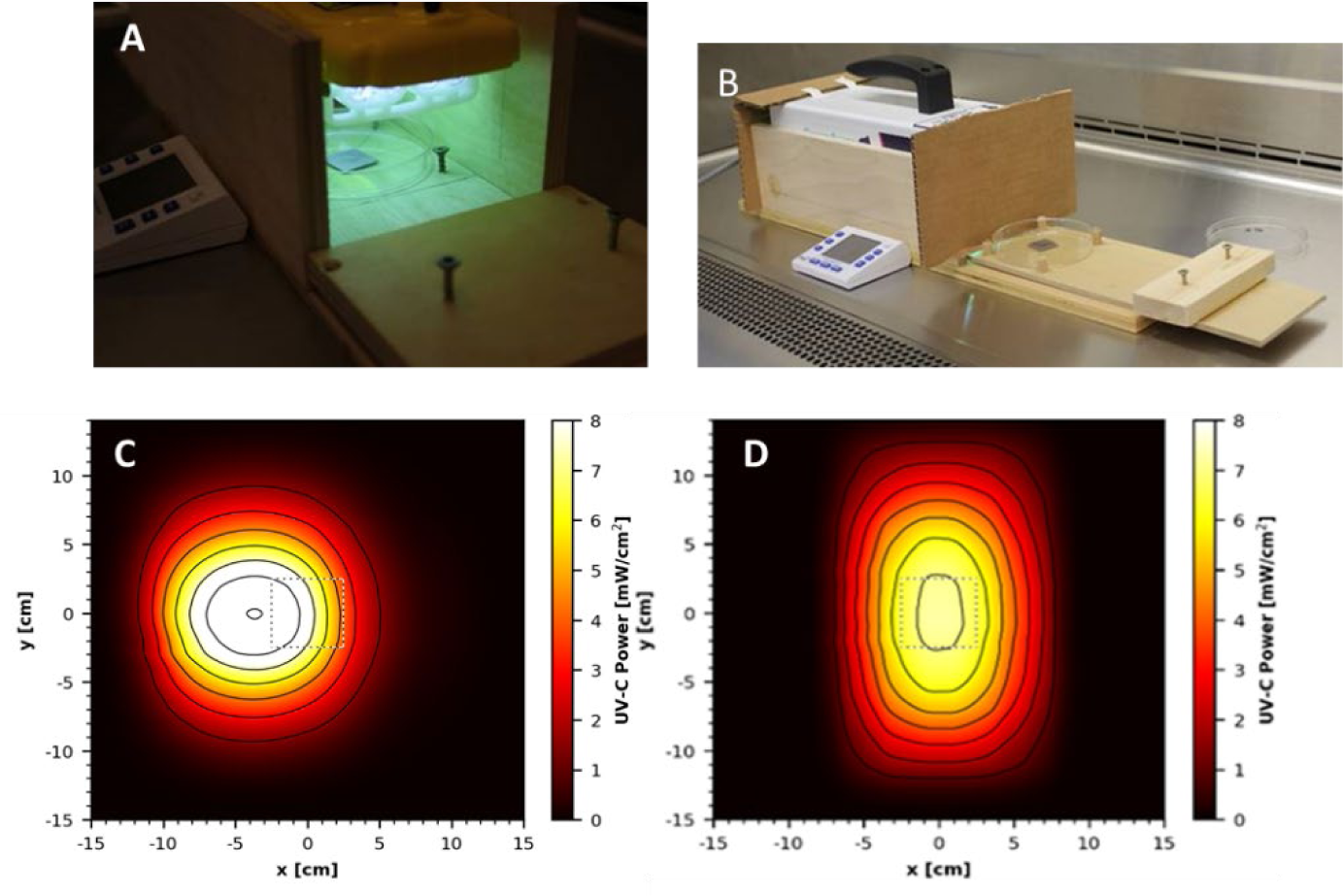
Testing setup and UV coverage for 18W and 35W handheld devices. **(A)** GermAwayUV 18W handheld device and custom test chamber, shown during a coupon exposure. **(B)** GermAwayUV 35W handheld device and custom test chamber, shown in the pre/post exposure state. **(C)** and **(D)** UV coverage heat maps for the 18W device **(C)** and the 35W device **(D)** taken 5 cm from the source.

For testing the two handheld devices, wooden holding chambers were constructed in which the devices could be placed to provide standardized exposures to test materials. They were designed to hold the UV source 5 cm above the surface of a test coupon, to prevent UV reflection, and to allow coupons to be inserted into the apparatus via a sliding tray for a specified time period of virus inactivation and then promptly removed (**Figure 3A and 3B**). The design of the chambers was the same for the two devices and only varied in size to accommodate the different dimensions of each device. The handheld devices had variable UV output immediately after turning them on. In order to generate consistent dosage for lab-to-lab testing, the devices were powered on 30 min prior to testing and remained powered on for the duration of the test.

Cost and schedule limitations prevented a statistical test among multiple handhelds from different manufacturing batches to assess the variability within and among devices and to determine a minimum warm-up time for end users. The variability highlighted an important gap, dosage monitoring in the field, which needs addressed for end users. To prevent potential contamination, the test chambers and devices were wiped down with pH6.8-adjusted bleach prior to being positioned inside a biosafety cabinet (BSC) for testing.

The sliding tray was constructed to hold a sterile Petri dish via guides and included a stop bar to ensure that the sample would be consistently positioned directly under the center of the UV source for maximum exposure. A cardboard barrier was placed over the opening of the chamber to prevent premature UV exposure onto test coupons when the materials were outside the test chamber. The plastic lid was removed from the Petri dish prior to UV exposure and the dish was wide enough that the dish edges did not impede UV transmission.

A N=5 was tested for each material at each time point. Each of the 5 coupons was inoculated with an independent virus preparation, emphasizing statistical accuracy over precision, and 3 separate exposures were tested for a total N=15. Test chambers held the UV source at a distance of 5 cm from the coupons, with the exception of keyboard keys. The keyboard keys were taller, and the distance from the UV bulb was 4.28-4.38 cm. The 18W and 35W handheld devices emitted steady state intensities of 10.12 mW cm^-2^ and 6.9 mW cm^-2^ respectively at the geometric center under the device. Test coupons were exposed to 10 or 20 seconds (s) of UV-C radiation from the 18W handheld and 2, 5, or 10 s of UV-C radiation from the 35W handheld. Different exposure times for the two devices were chosen based on pre-experimental predictions that were considered for practical application of the devices in a field setting. Prior to testing it was assumed that 35W radiation would exceed 18W and 10 s was a common time variable for both the 18W and 35W handhelds. During testing, the ambient environment was 22±2°C and 40% RH. The surface temperature within the test chamber reached 36°C under the 18W device and 48°C under the 35W device. Following UV exposure, coupons were transferred using sterile forceps to 50-ml conical tubes for extraction. The corresponding virus-inoculated control coupons were left at ambient conditions during testing because prior testing with Φ6 showed complete recovery/survival of >8 log_10_ of dried virus after treatment of different surfaces at 55°C, 50% RH for 1 h (Buhr et al. 2020). Of note, wet virus was completely killed at 55°C, 50% RH for 1 h (Buhr et al. 2020), an important data point that supported the goal for testing dried virus as described in the introduction and discussion. Hence, the heat (48°C) could have potentially impacted decontamination kinetics if the tests had used wet virus instead of dried virus. The UV test parameters here were short duration (up to 10 s) and the surface temperatures likely did not equilibrate much above ambient temperature. Importantly, the high temperatures measured under these handhelds generates a practical risk for the end users because it is not known if the heat generated might create a fire hazard if the lamps are left on for extended periods. While 48°C was the highest temperature measured among any of the devices, temperature measurements over extended run periods for multiple different handheld batches would be needed for safety assessments prior to field use, and this task was outside of the testing scope.

The dosage and virus inactivation results are summarized in **Tables 2** (log_10_ reduction) and **3** (log_10_ survival). Dosages and virus inactivation were measured at a 5 cm distance, which was considered a reasonable, practical distance for a handheld device used to scan over surfaces. The keyboard keys were slightly taller and closer to the UV source. Thus, the dosage on the keys was slightly greater than the other materials but no dosage calculations were made specifically for those keys.

**Table 2.**
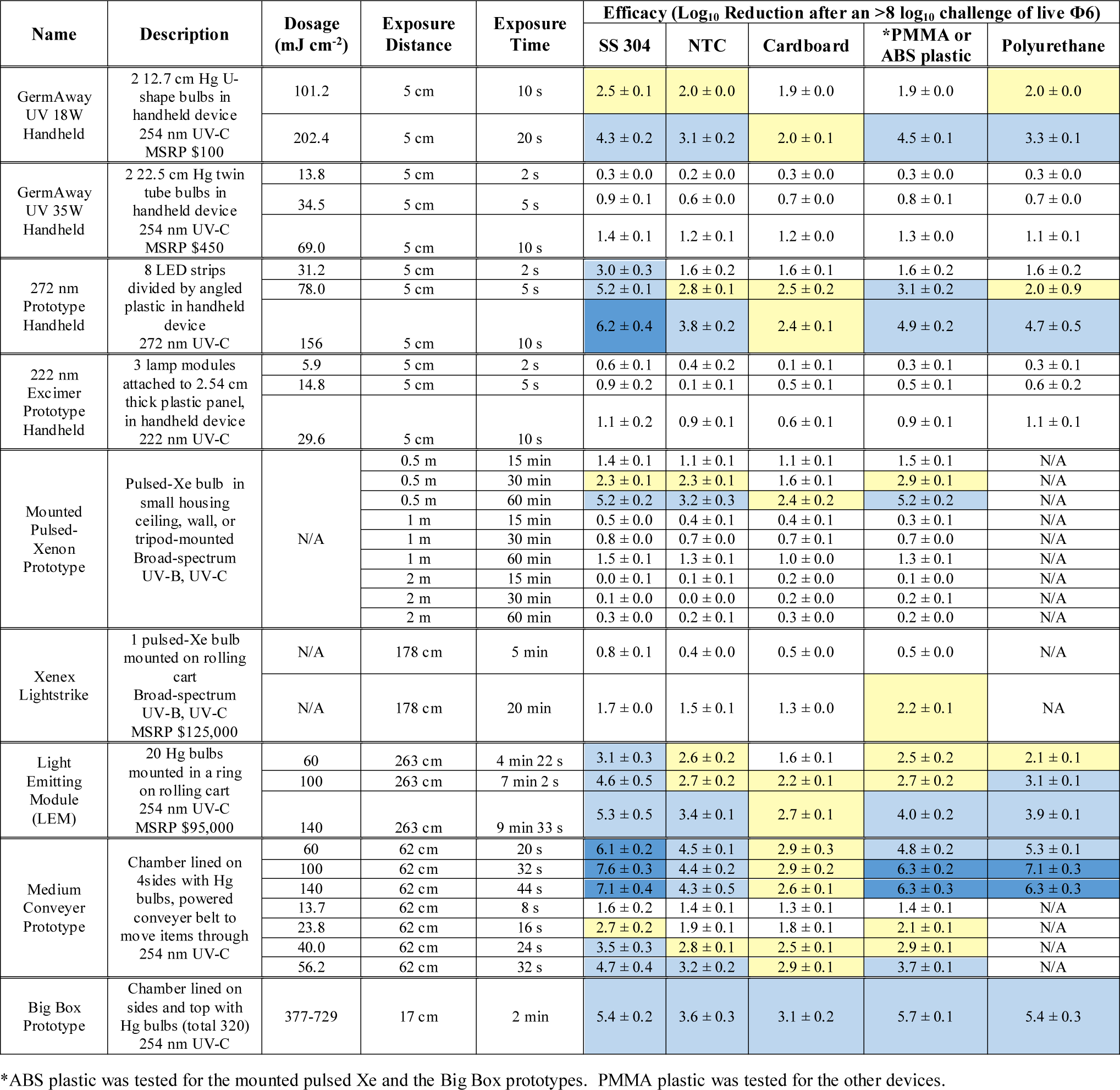
Dosage and efficacy of handheld, room, and chamber-type devices showing log_10_ reduction data based on the steady state emission, not peak emission. Legend: White (low decontamination) = Fail <2 log_10_; Yellow (sanitation) = Fail ≥2 log_10_, <3 log_10_; Light Blue (disinfection) = Pass ≥ 3 log_10_; and Dark Blue (approaching virus sterilization) = Pass ≥ 6 log_10_. N/A – dosage measurements had no meaning because of the broad-spectrum Xe source.

To evaluate the efficacy of the devices, a minimum of 3-log_10_ inactivation was targeted, which is equivalent to a 99.9% reduction and corresponds to the current EPA requirements for chemical disinfection. A 10 s exposure with the GermAway 18W unit failed to meet the ≥3 log_10_ inactivation threshold for all tested materials. A 20 s exposure successfully achieved a greater than 3 log_10_ inactivation out of an 8.2 log_10_ virus challenge on SS304, NTC, keyboard keys, and polyurethane but failed to meet the 3 log_10_ inactivation threshold on cardboard.

The GermAwayUV 35W handheld sanitizer failed to meet the ≥3 log_10_ inactivation threshold out of an 8 log_10_ PFU virus challenge on all five materials for all three exposure durations, achieving less than 2 log_10_ PFU inactivation. The GermAwayUV 35W handheld sanitizer delivered lower dosage than the 18W handheld despite nearly double the power. Hence there was no correlation between power and dosage/efficacy, and the importance of measuring every device was apparent.

### Prototype Handheld devices (272 nm LED and 222 nm Lamp Modules)

Two additional handheld devices were tested for efficacy of virus inactivation, which were prototypes rather than commercial units. The first prototype was one of two custom 3-D printed proprietary units and featured eight LED strips which emitted 272 nm wavelength UV-C radiation. The face of the handheld was 320 mm x 100 mm with the LED strips covering 255 mm x 60 mm. An average intensity of 12.71 mW cm^-2^ was measured within a decontamination footprint of 6 cm x 25.5 cm at 5 cm standoff distance from the bulb (**Figure 4A and 4B**). The second prototype device utilized three 222 nm UV-C Excimer Lamp Modules installed into a 2.54 cm thick white plastic panel with power supply. It is important to note that this was strictly an early prototype undergoing iterative improvements, and the UV sources were spaced too far apart for a wand configuration. An average intensity of 1.54 mW cm^-2^ was measured at 5 cm standoff distance from an individual module (**Figure 5A and 5B**).

**Figure 4.**
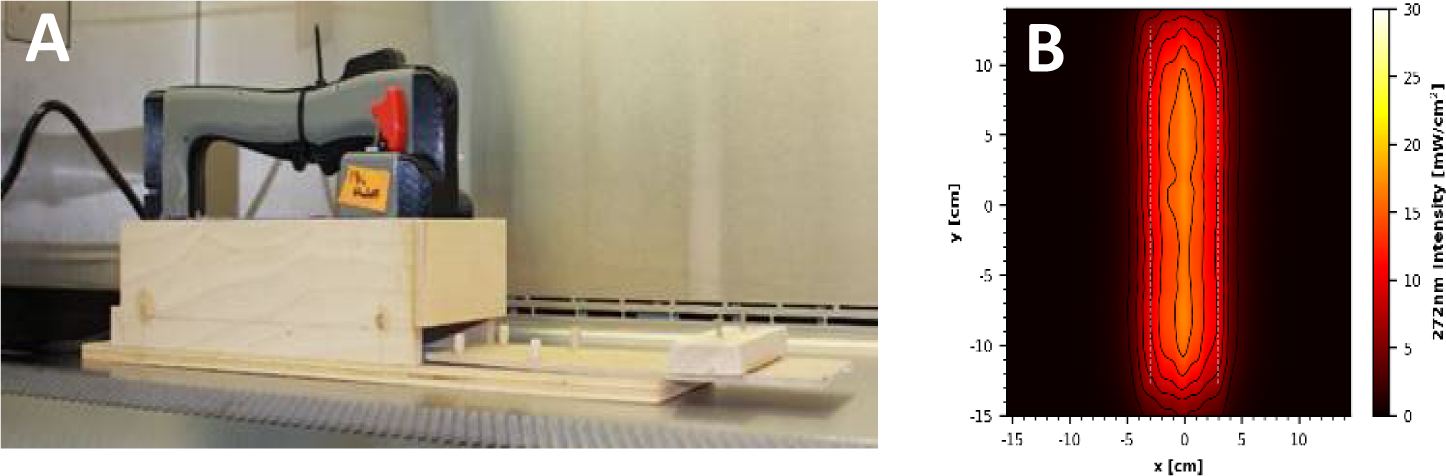
**(A)** Prototype 272 nm LED handheld inside wooden test chamber. **(B)** UV coverage heat map taken 5 cm from the source.

**Figure 5.**
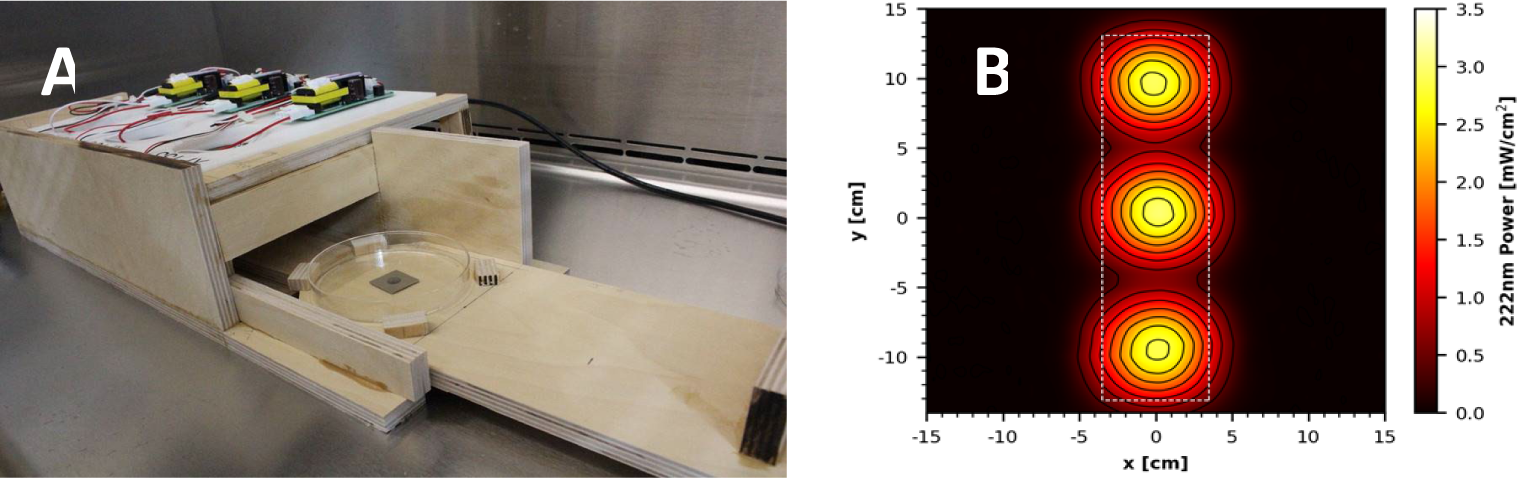
**(A)** Prototype 222 nm Excimer Lamp Module Board inside wooden test chamber. **(B)** UV coverage heat map taken 5 cm from the source.

The wooden test chambers for the prototype handhelds followed the same design as those for the 18W and 35W devices, with the additional feature of a wooden barrier that removed the need for cardboard to prevent premature UV exposure onto test coupons when the materials were outside the UV chamber. Again, there was a 5 cm vertical standoff distance from the UV source to the surface of the test coupons. Mimicking the 18W and 35W handheld unit tests, the devices were powered on 30 min prior to testing to warm up and remained powered on for the duration of the test. An Ophir Spiricon Starbright Dosimeter (S/N 949685, P/N 7201580) and sensor (S/N 954282, P/N 7Z02479) were used to confirm that the 222 nm device was on and emitting 222 nm UV radiation, as the design of the prototype did not allow visual confirmation that the device was on after it was plugged in. The test chambers and handheld UV devices were wiped down with pH6.8-adjusted bleach prior to being positioned inside a BSC for testing.

A N=5 coupons for each material were tested at each time/dosage with each coupon inoculated with an independent virus preparation. During tests, virus-inoculated coupons were transferred singly to sterile Petri plates and inserted into the test chambers via the sliding tray for timed UV exposures at the geometric center of the handheld device. For the 272 nm device, the cardboard coupons were anchored down using sterile pipette tips due to the large amount of air movement generated by the cooling fans of the device. In the 272 nm prototype, coupons were exposed to a steady state intensity of 15.6 mW cm^-2^ measured at the geometric center of the device with a 5 cm standoff distance. Similarly, the 222 nm prototype emitted an intensity of 2.96 mW cm^-2^ at a similar location centered under a single lamp module. Following UV exposure, the coupons were transferred to 50-ml conical tubes for extraction. For both devices, test coupons were exposed to UV-C radiation for 2, 5, or 10 s. For the 272 nm device, the ambient environment during testing was 21±2°C and 21% RH, and the surface temperature under the sterilizer reached 34.7±2°C. For the 222 nm device, the ambient environment was 21.8±2°C and 20% RH, and the surface temperature reached 28.3±2°C within the test chamber.

The dosage and virus inactivation results are summarized in **Tables 2** (log_10_ reduction) and **3** (log_10_ survival). The 272 nm LED prototype successfully achieved a ≥3 log_10_ PFU inactivation out of an 8.5 log_10_ PFU virus challenge for SS304 at 2, 5, and 10 s, for ABS at 5 and 10 s, and for NTC and polyurethane at 10 s. The hardest, smoothest material was SS304, and it showed the greatest log_10_ reduction at all 3 time points. Cardboard showed the lowest inactivation rate with no treatments providing ≥3 log_10_ PFU inactivation. Overall, the 272 nm LED prototype showed significantly greater virus inactivation compared to the 18W and 35W handheld commercial devices.

The 222 nm Excimer UV prototype failed to achieve a >3 log_10_ inactivation out of an 8.5 log_10_ virus challenge for all 5 materials tested, making it the least effective of the four handheld devices tested. Further testing with longer exposure times might produce results passing the ≥3 log_10_ inactivation threshold. From a practical standpoint, this data showed that this 222 nm prototype had poor efficacy and very limited utility. Since this was a prototype, iterative improvements can be made to improve performance of this device.

### Prototype Mounted Pulsed Xenon Unit for Room Decontamination

A prototype room-decontamination unit featuring a pulsed Xe UV bulb was tested. The unit consists of a pulsed Xe bulb within a frame intended to be mounted onto a wall, ceiling, or mobile tripod for room decontamination. The UV source emitted a small burst of broad-spectrum radiation every 6 s with the burst lasting for a short duration. The electromagnetic spectrum included UV-C, UV-B, UV-A, and violet-blue visible radiation. Reflector material was positioned behind the source to enhance UV output.

Testing of the modified prototype took place within an enclosure provided by the vendor.

The device was mounted at 0.5, 1, and 2 m vertical standoff distance above the testing surface (**Fig. 6**). Test coupons were placed below the prototype in sterile petri dishes and aseptic technique was employed to the greatest extent possible while outside of a BSC to prevent contamination. The coupons contained within Petri dishes were uncovered just prior to the test and re-covered at the conclusion of the exposure times. Independent tests were run for 3 exposure times (15, 30, and 60 min), each taken at 0.5, 1, and 2 m distances from the UV source. These time increments were determined via the recommended cycle lengths from the vendor and corresponded to vendor test data (30 and 60 min only). The device was pre-programmed for 30 min run times, therefore for the 15-min increment, coupons were removed from the enclosure without shutting off the device after 15 min had elapsed from the time of the first flash. For the 60-min cycle, two decontamination cycles were run sequentially.

**Fig. 6.**
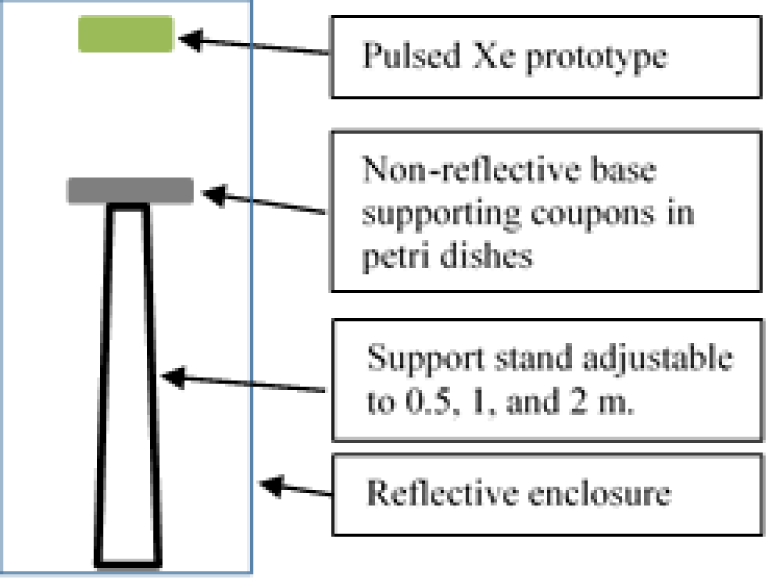
Virus-inoculated coupons were placed 0.5, 1, or 2 m below the Pulsed Xe prototype.

Results for a mounted prototype containing a pulsed Xe bulb are shown in **Tables 2** (log_10_ reduction) and **3** (log_10_ survival). This device emitted broad-spectrum radiation in pulses occurring every 6 s with the duration of each pulse measured at 0.489 s, and the majority of the dosage applied over the first few milliseconds of that time. Because of the broad spectrum nature, the UV dosage could not be confidently measured. This device demonstrated measurable efficacy at 0.5 m for 60 min, and the results were best on non-porous materials. Efficacy was very limited at 1 and 2 m and shorter exposure times, particularly on porous cardboard, followed by semi-porous NTC. As usual, the best efficacy was on the smooth surfaces: plastic and SS304.

### Commercial Rolling Units for Room Decontamination

Two commercial rolling units designed for room decontamination were purchased. The first was the Xenex Lightstrike (Model PXUV4D, S/N 002628, Xenex Disinfection Systems, San Antonio, TX, USA), which contained one pulsed Xe bulb (broad spectrum across the germicidal spectrum of 200-315 nm). The bulb extends and retracts at the top of the unit and pulsed at a rate of 67 flashes per s. The intensities and dosages at specific wavelengths were not carefully analyzed/dissected because the work was not aimed at correlating specific wavelength dosages from a broad spectrum device to a kill rate. The second unit was the Light Emitting Module (“LEM,” Rapid UV-C Disinfection Model R3, S/N 473, 120V/12A, STERILIZ, LLC, 150 Linden Oaks, Rochester, NY 14625-2802), which contained a ring of twenty Hg bulbs with a 41- cm diameter that emitted predominantly 254 nm wavelength UV-C radiation. The device was tested at an exposure distance of 2.63 m from the center of the Hg bulb ring (**Fig. 7**). The length of exposure was controlled based upon the cumulative dosage recorded via the LEM system dosimeters placed next to the test coupons and targeted for exposures of 60, 100, and 140 mJ cm^-2^. Coupons were exposed to an average intensity calculated to be 0.23-0.24 mW cm^-2^. Due to the different expected intensities of the UV sources, the devices were set at different distances from test coupons to achieve similar dosages in an attempt to directly compare the killing efficacy of a broad spectrum radiation source to a 254 nm source.

**Fig. 7.**
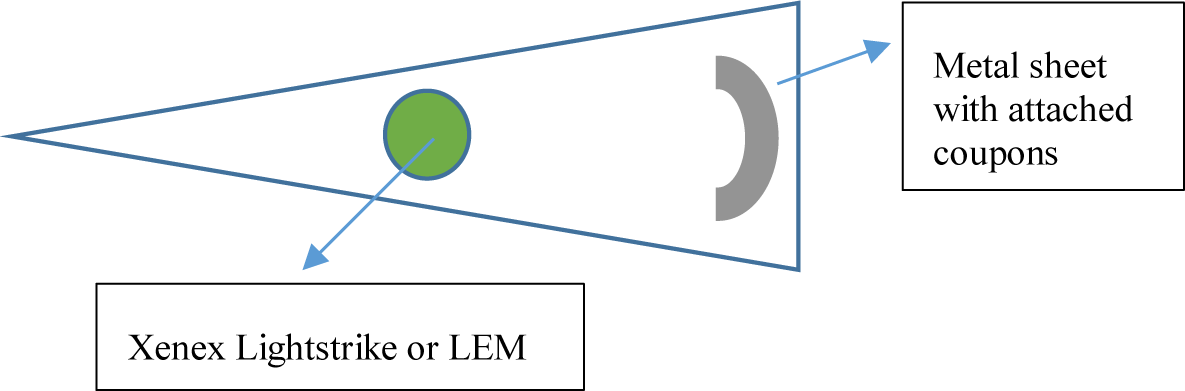
Virus-inoculated coupons were placed 1.78 m horizontally from the Xenex Lightstrike or 2.63 m away from the LEM.

For testing, the Xe or Hg rolling units were positioned in the corner of a triangular area and non-reflective folding panels were set up to prevent UV exposure to personnel outside of the decontamination area. Magnets were glued to the underside of test coupons prior to inoculation of virus, and a black, non-reflective, metal sheet rack was utilized as a support for the test coupons. The rack was bent into a curved shape in an attempt to maintain a constant UV exposure distance to all coupons. Testing of these two devices required transport of coupons to the testing site, and coupons were transported in 50-ml conical tubes at room temperature.

Negative control coupons as well as additional shipping controls (inoculated and transported, but not exposed to UV) were also included. Conditions in the testing room were not aseptic but care was taken to avoid contamination at each step and coupons were only transferred to and from the metal rack using sterile forceps. After UV exposure, samples were transferred to new sterile conical tubes and transported back to the microbiology lab for virus extraction and quantification.

Specific testing conditions differed slightly between the two rolling units. For testing the Xenex Lightstrike, the metal stand holding virus-inoculated test coupons was placed such that the coupon height was between 1.09 and 1.55 m above the ground (approximately parallel to the height of the pulsed Xe bulb), and the distance between the coupons and the UV source was 1.72-1.78 m. Based on preliminary dosage readings, the Xenex Lightstrike did not need a 30-min warm up time. Two time points of 5 and 20 min were tested. Room conditions were measured at 23.3±1°C and 74% RH for the first exposure and 25.1±1°C and 22% RH for the second exposure. As the tests occurred in succession approximately 30 min apart, the shift in environmental conditions with the rise in temperature and drop in humidity is speculated to be driven by the Xe unit itself. Additionally, the smell of ozone was detected in the air following the completion of each test. Ozone level in the room was measured at 0.26 ppm for the Xenex Lightstrike (0.1 ppm is the 8-h Occupational Safety and Health Assessment (OSHA) limit).

For testing the LEM, the metal stand holding virus-inoculated test coupons was placed such that the coupon height was approximately 1.2 m above the ground (parallel to center of the Hg bulbs), and the distance between the center of the ring of UV bulbs to the center of the metal arc with coupons was approximately 2.62 m. The distance between the test coupons and the nearest UV bulb was 2.43 m. Testing for this device included three independent exposures of 60, 100, and 140 mJ cm^-2^ which took 4 min 22 s, 7 min 2 s, and 9 min 33 s, respectively. The exposure conditions were 26±1°C, 38% RH. Ozone level in the room was measured at 0.08 ppm for the LEM (0.1 ppm is the 8-h Occupational Safety and Health Assessment (OSHA) limit).

Results for the Xenex Lightstrike unit with pulsed Xe UV bulb are shown in **Tables 2** and **3**. The Xenex Lightstrike failed to achieve a ≥3 log_10_ inactivation out of an 8.4 log_10_ PFU virus challenge for all 5 materials tested. A positive is that the Xenex did not require a warmup time in contrast to devices with Hg bulbs.

**Table 3.** Dosage and efficacy of handheld, room, and chamber-type devices showing log_10_ survival data based on the steady state emission, not peak emission. Legend: White (low decontamination) = Fail <2 log_10_; Yellow (sanitation) = Fail ≥2 log_10_, <3 log_10_; Light Blue (disinfection) = Pass ≥ 3 log_10_; and Dark Blue (approaching virus sterilization) = Pass ≥ 6 log_10_. N/A – dosage measurements had no meaning because of the broad-spectrum Xe source.

| Name | Description | Dosage (mJ cm<sup>-2</sup>) | Exposure Distance | Exposure Time | Log<sub>10</sub> Survival after a >8 log<sub>10</sub> challenge of live Φ6 |  |  |  |  |  |
| --- | --- | --- | --- | --- | --- | --- | --- | --- | --- | --- |
|  |  |  |  |  | Inoculum | SS 304 | NTC | Cardboard | *PMMA or ABS plastic | Polyurethane |
| GermAway UV 18W Handheld | 2 12.7 cm U-shape Hg bulbs in handheld device 254 nm UV-C MSRP \$100 | 101.2 | 5 cm | 10 s | 8.4 +/- 0.1 | 5.9 ± 0.5 | 6.4 ± 0.1 | 6.5 ± 0.1 | 5.5 ± 1.1 | 6.4 ± 0.1 |
|  |  | 202.4 | 5 cm | 20 s | 8.2 +/- 0.0 | 3.9 ± 0.4 | 5.0 ± 0.4 | 6.1 ± 0.3 | 3.7 ± 0.3 | 5.1 ± 0.2 |
| GermAway UV 35W Handheld | 2 22.5 cm twin tube Hg bulbs in handheld device 254 nm UV-C MSRP \$450 | 13.8 | 5 cm | 2 s | 8.2 +/- 0.1 | 7.8 ± 0.2 | 7.9 ± 0.1 | 7.8 ± 0.1 | 7.8 ± 0.2 | 7.8 ± 0.1 |
|  |  | 34.5 | 5 cm | 5 s |  | 7.2 ± 0.2 | 7.5 ± 0.1 | 7.3 ± 0.1 | 7.3 ± 0.3 | 7.4 ± 0.1 |
|  |  | 69.0 | 5 cm | 10 s |  | 6.6 ± 0.3 | 6.9 ± 0.2 | 6.9 ± 0.1 | 6.8 ± 0.2 | 7.0 ± 0.2 |
| 272 nm Prototype Handheld | 8 LED strips divided by angled plastic in handheld device 272 nm UV-C | 31.2 | 5 cm | 2 s | 8.5 +/- 0.2 | 5.5 ± 0.6 | 7.0 ± 0.3 | 6.9 ± 0.2 | 6.9 ± 0.3 | 6.9 ± 0.4 |
|  |  | 78.0 | 5 cm | 5 s |  | 3.3 ± 0.0 | 5.7 ± 0.1 | 6.0 ± 0.4 | 5.4 ± 0.3 | 6.5 ± 1.9 |
|  |  | 156 | 5 cm | 10 s |  | 2.3 ± 0.7 | 4.7 ± 0.3 | 6.1 ± 0.1 | 3.6 ± 0.5 | 3.8 ± 0.9 |
| 222 nm Excimer Prototype Handheld | 3 lamp modules attached to 2.54 cm thick plastic panel, in handheld device 222 nm UV-C | 5.9 | 5 cm | 2 s | 8.3 +/- 0.2 | 7.7 ± 0.2 | 7.9 ± 0.3 | 8.2 ± 0.1 | 8.0 ± 0.1 | 7.9 ± 0.0 |
|  |  | 14.8 | 5 cm | 5 s |  | 7.4 ± 0.3 | 8.2 ± 0.1 | 7.8 ± 0.1 | 7.8 ± 0.2 | 7.6 ± 0.3 |
|  |  | 29.6 | 5 cm | 10 s |  | 7.1 ± 0.4 | 7.1 ± 0.4 | 7.6 ± 0.1 | 7.4 ± 0.2 | 7.1 ± 0.1 |
| Mounted Pulsed-Xenon Prototype | Pulsed-Xe bulb in small housing ceiling, wall, or tripod-mounted Broad-spectrum UV-B, UV-C | N/A | 0.5 m | 15 min | 8.2 +/- 0.1 | 6.8 ± 0.2 | 7.0 ± 0.2 | 7.1 ± 0.2 | 6.7 ± 0.1 | N/A |
|  |  |  | 0.5 m | 30 min |  | 5.9 ± 0.3 | 5.8 ± 0.3 | 6.6 ± 0.1 | 5.3 ± 0.2 | N/A |
|  |  |  | 0.5 m | 60 min |  | 2.9 ± 0.4 | 5.0 ± 0.8 | 5.7 ± 0.5 | 3.0 ± 0.6 | N/A |
|  |  |  | 1 m | 15 min |  | 7.7 ± 0.1 | 7.7 ± 0.2 | 7.7 ± 0.1 | 7.8 ± 0.1 | N/A |
|  |  |  | 1 m | 30 min |  | 7.4 ± 0.1 | 7.5 ± 0.0 | 7.4 ± 0.1 | 7.4 ± 0.1 | N/A |
|  |  |  | 1 m | 60 min |  | 6.7 ± 0.2 | 6.8 ± 0.2 | 7.2 ± 0.1 | 6.8 ± 0.2 | N/A |
|  |  |  | 2 m | 15 min |  | 8.2 ± 0.1 | 8.1 ± 0.1 | 8.0 ± 0.1 | 8.1 ± 0.0 | N/A |
|  |  |  | 2 m | 30 min |  | 8.1 ± 0.1 | 8.1 ± 0.1 | 8.0 ± 0.1 | 8.0 ± 0.1 | N/A |
|  |  |  | 2 m | 60 min |  | 7.9 ± 0.0 | 7.9 ± 0.1 | 7.8 ± 0.1 | 7.9 ± 0.1 | N/A |
| Xenex Lightstrike | 1 pulsed-Xe bulb mounted on rolling cart Broad-spectrum UV-B, UV-C MSRP \$125,000 | N/A | 178 cm | 5 min | 8.4 +/- 0.0 | 7.6 ± 0.2 | 8.0 ± 0.1 | 7.9 ± 0.1 | 7.8 ± 0.1 | N/A |
|  |  | N/A | 178 cm | 20 min |  | 6.7 ± 0 | 6.9 ± 0.3 | 7.1 ± 0.0 | 6.2 ± 0.1 | N/A |
| Light Emitting Module (LEM) | 20 Hg bulbs mounted in a ring on rolling cart 254 nm UV-C MSRP \$95,000 | 60 | 263 cm | 4 min 22 s | 8.4 +/- 0.2 | 5.3 ± 0.6 | 5.8 ± 0.5 | 6.8 ± 0.2 | 5.9 ± 0.5 | 6.3 ± 0.1 |
|  |  | 100 | 263 cm | 7 min 2 s |  | 3.8 ± 1.1 | 5.7 ± 0.5 | 6.2 ± 0.2 | 5.7 ± 0.4 | 5.3 ± 0.1 |
|  |  | 140 | 263 cm | 9 min 33 s |  | 3.1 ± 1.1 | 5.0 ± 0.3 | 5.8 ± 0.2 | 4.4 ± 0.3 | 4.6 ± 0.2 |
| Medium Conveyor Prototype | Chamber lined on 4 sides with Hg bulbs, powered conveyor belt to move items through 254 nm UV-C | 60 | 62 cm | 20 s | 8.2 +/- 0.1 | 2.1 ± 0.4 | 3.7 ± 0 | 5.3 ± 0.6 | 3.4 ± 0.4 | 2.9 ± 0.1 |
|  |  | 100 | 62 cm | 32 s |  | 0.6 ± 0.6 | 3.8 ± 0.5 | 5.3 ± 0.4 | 2.0 ± 0.4 | 1.1 ± 0.7 |
|  |  | 140 | 62 cm | 44 s |  | 1.1 ± 0.8 | 3.9 ± 1.1 | 5.6 ± 0.1 | 1.9 ± 0.6 | 0.9 ± 0.5 |
|  |  | 13.7 | 62 cm | 8 s | 8.4 +/- 0.2 | 6.8 ± 0.4 | 7.0 ± 0.1 | 7.1 ± 0.1 | 7.0 ± 0.1 | N/A |
|  |  | 23.8 | 62 cm | 16 s |  | 5.6 ± 0.4 | 6.5 ± 0.1 | 6.5 ± 0.2 | 6.3 ± 0.1 | N/A |
|  |  | 40.0 | 62 cm | 24 s |  | 4.8 ± 0.6 | 5.5 ± 0.2 | 5.8 ± 0.3 | 5.4 ± 0.2 | N/A |
|  |  | 56.2 | 62 cm | 32 s |  | 3.7 ± 1.0 | 5.1 ± 0.4 | 5.5 ± 0.2 | 4.7 ± 0.1 | N/A |
| Big Box Prototype | Chamber lined on sides and top with Hg bulbs (total 320) 254 nm UV-C | 377-729 | 17 cm | 2 min | 8.4 +/- 0.1 | 3.0 ± 0.4 | 4.8 ± 0.6 | 5.3 ± 0.5 | 2.5 ± 0.3 | 3.0 ± 0.6 |
|  |  |  |  |  | 8.2 +/- 0.1 (ABS plastic only) |  |  |  |  |  |

\*ABS plastic was tested for the mounted pulsed Xe and the Big Box prototypes. PMMA plastic was tested for the other devices.

Results for the LEM with Hg bulbs are shown in **Tables 2** and **3**. The LEM successfully achieved a ≥3 log_10_ PFU inactivation out of an 8.4 log_10_ PFU virus challenge for SS304 at all three dosages, for polyurethane at the higher two exposures, and for NTC and keyboard keys at the highest dosage only. It failed to meet the ≥3 log_10_ PFU inactivation threshold for cardboard at all three exposure levels.

### Prototype Medium Conveyer

The prototype medium conveyer featured a chamber measuring 2.03 m long x 0.78 m wide x 0.69 m tall that was lined on all interior surfaces with UV-C emitting (254 nm) Hg bulbs, including below the powered rollers (**Figure 8**). Testing of this device required transport of coupons to the testing site, and coupons were transported in 50-ml conical tubes at room temperature. Negative control coupons as well as additional shipping controls (inoculated and transported, but not exposed to UV) were also included.

**Figure 8.**
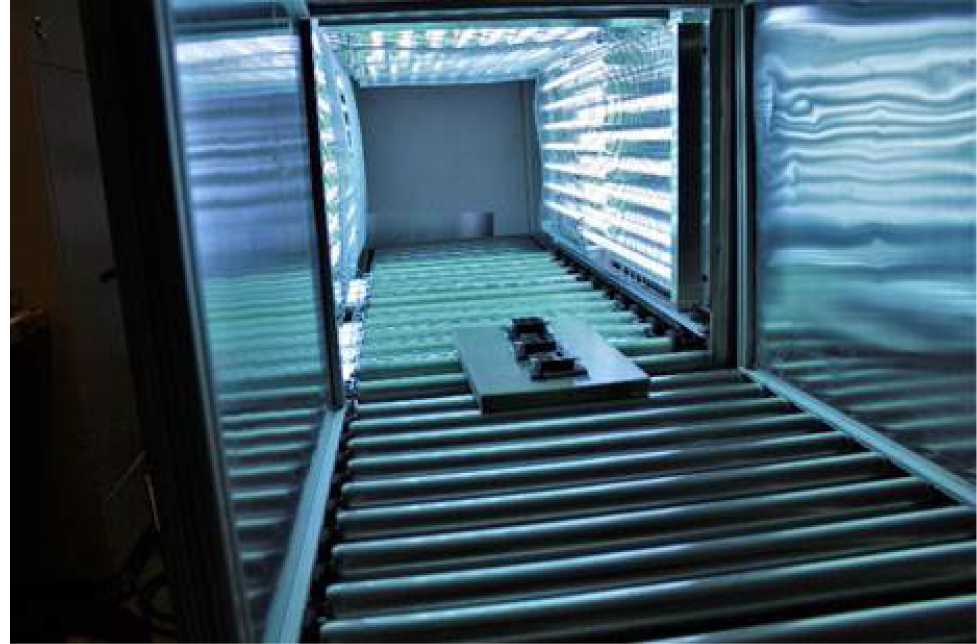
Dosimeters traveling down conveyor to determine exposure times for target UV dosages.

Two rounds of testing were performed with the conveyer device, with slight differences in experimental setup and UV dosages. For both rounds, dosimeters were first used in trial-and-error runs to determine the required run-through time to reach the target UV exposures. The dosimeters used were Roithner LaserTechnick GmbH GIVA-S12SD dosimeters from Vienna, Austria, with dimensions of 4.3 cm x 3.5 cm x 1.8 cm. In the first round of experiments, three dosimeters were horizontally taped to a 2% polyethylene board (46.7 cm x 28.6 cm x 2.54 cm) and were sent through the conveyor to get dosage readings based on exposure time (**Figure 8**). After target exposure times were determined, coupons were placed inside sterile Petri dishes and set on the same polyethylene support board before exposure in the conveyer. The first round of testing included exposures of 60 mJ cm^-2^ (22 s), 100 mJ cm^-2^ (32 s), and 140 mJ cm^-2^ (44 s). Conditions within the conveyer for this round were 27.7 °C, 63.2% RH, 0.09 ppm ozone.

In the second round of testing, the initial runs were again dosimeter-only to determine exposure times to reach the targeted UV dosages. The same dosimeters were used, but this time they were placed on a ceramic tile (∼45.7 cm x 45.7 cm). During testing, coupons were placed directly on the ceramic tile support to prevent the sides of the Petri dishes from blocking any UV radiation from reaching the coupons. Test conditions were 17.3±1°C and 20.1% RH. Ozone reading was not captured since the ozone reader was unavailable.

Two rounds of testing were carried out for the prototype medium conveyer with Hg bulbs, with each round varying in dosages tested and in the method of exposing the test coupons. The dosages over time were not perfectly linear. The dosage variability over time might have been variability in dosimeter readings and/or variability in Hg bulb dosages after warmup. Test results are shown in **Tables 2** and **3**. During round 1 testing at 60 mJ cm^-2^ (20 s), 100 mJ cm^-2^ (32 s), and 140 mJ cm^-2^ (44 s), the conveyer successfully achieved a ≥3 log_10_ PFU inactivation out of an 8.2 log_10_ PFU virus challenge for all three exposure times on SS304, NTC, ABS plastic, and polyurethane, with slightly higher inactivation results for ABS plastic and polyurethane at the higher two treatments. It failed to meet the ≥3 log_10_ PFU inactivation threshold on cardboard for all three exposure times.

For round 2 of testing, the dosages measured during testing were 13.7 mJ cm^-2^ (8s), 23.8 mJ cm^-2^ (16s), 40.0 mJ cm^-2^ (24s), and 56.2 mJ cm^-2^ (32s). Regardless of dosage variability, the conveyer successfully achieved a ≥3 log_10_ inactivation out of an 8.4 log_10_ virus challenge for SS304 at 24 and 32 s, for NTC at 32 s, and for ABS plastic at 32 s time points. For all other materials and round 2 exposure times, it failed to reach the ≥3 log_10_ PFU threshold inactivation.

The conveyor produced UV dose-dependent inactivation at lower dosages (13.7-56.2 mJ cm^-2^), but inactivation leveled off across all surfaces tested at higher dosages (60-140 mJ cm^-2^). The size of the shielded virus population was dependent on material porosity since the highest level of inactivation was observed on non-porous SS304, followed by polyurethane, ABS plastic, NTC, and then porous cardboard. In addition, a sub-population of virus protected by debris was shielded from exposure to radiation because of the presence of host cell debris as indicated by a flattening of the kill rate across all the materials including smooth SS304. That sub-population of debris-complexed virus manifest may manifest higher resistance to the damaging effects of the UV radiation because of both shielding and drying; it is widely known that UV damage produces pyrimidine dimers in nucleic acid and biochemical reactions involving bond formation typically require a solvent like water.

### Prototype Big Box UV Chamber for Pallets

A prototype Big Box UV Sterilizer, a proprietary UV-C decontamination device, was acquired for virus inactivation testing. The outside dimensions were 2.74 m x 2.24 m x 2.4 m with an interior large enough to accommodate a recommended maximum load with dimensions of 1.21 m wide x 1.21 m long x 1.52 m tall. Max interior load was 1,134 kg. The interior was lined on five surfaces with a total of 320 T8 Hg bulbs, each measuring 0.9 m long and emitting 254 nm UV-C radiation. A double-stacked pallet mock-up of dimensions 1.02 m x 1.22 m x 1.64 m was placed within the UV chamber (**Figure 9**), centered from left to right, and positioned up against the rear backstop on the base of the chamber.

**Figure 9.**
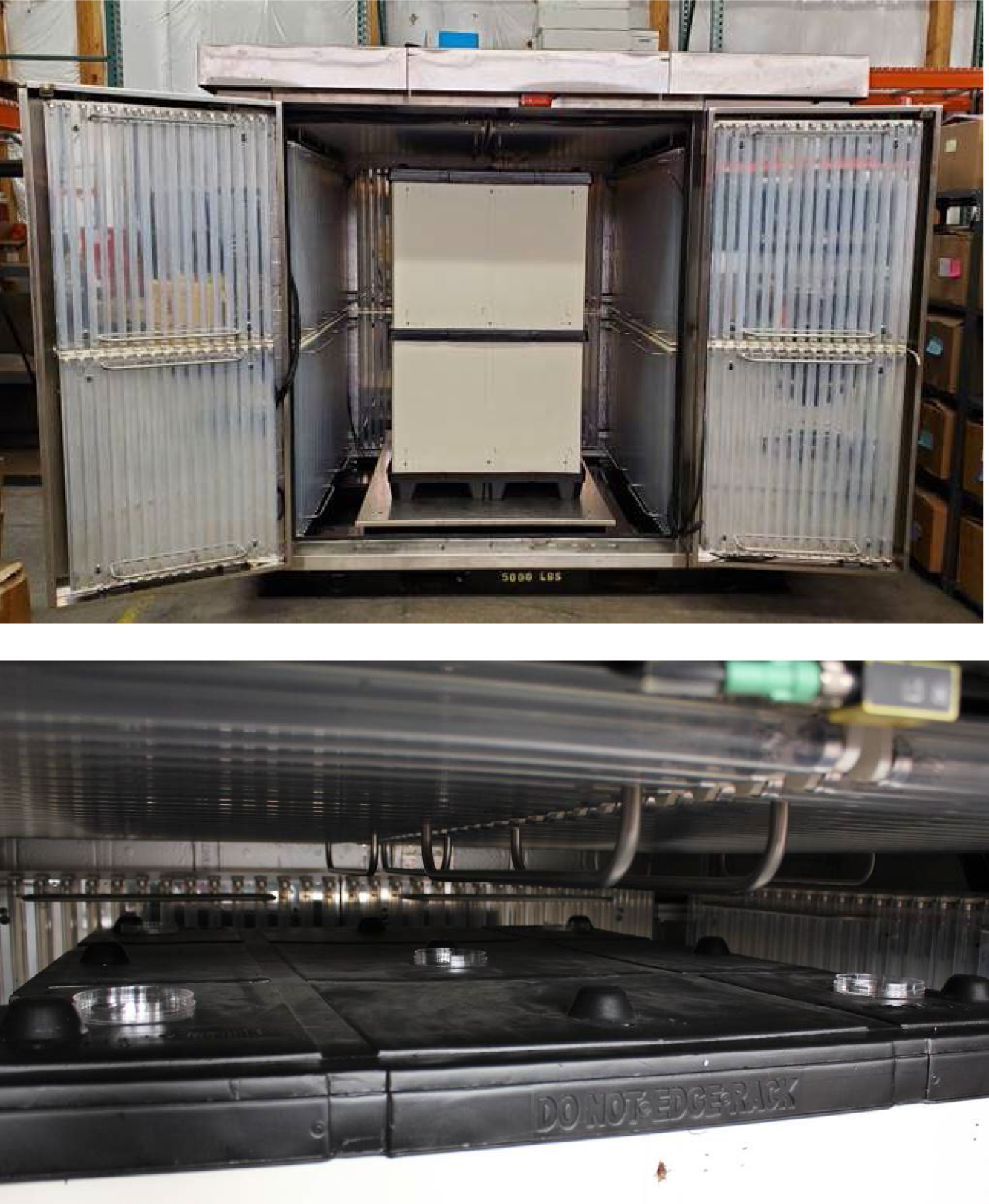
Prototype Big Box UV-C Chamber with a double stacked pallet (top). Coupon placement inside petri plates on top of pallet for testing (bottom).

During testing, coupons were placed in Petri dishes on top of the pallet in five separate locations with lids removed prior to exposure. The UV chamber doors were closed, and the chamber was operated via a pre-programmed cycle set to run for 2 min followed by a 30 s exhaust. After UV exposure, the coupons were recovered, and the surviving virus was extracted and quantified. There was a single combined 2 min exposure test run for all coupons except for ABS plastic coupons, which was tested for 2 min on a separate test day. Room temperature extraction control samples were transported to and from the test location along with test coupons. Peak ozone generated was 0.36 ppm. Ozone was purged out of the chamber top for 30 s prior to opening the doors.

Results for the prototype Big Box UV sterilizer are shown in summary **Tables 2** and **3**. A large double-stacked pallet mock-up was set inside the Big Box UV sterilizer. Coupons were then set on top of the plastic and cardboard mock-up for UV exposure, and the distance from virus-inoculated coupon to the nearest Hg bulbs on the chamber ceiling was 16.5 cm. The dosages varied significantly at different locations in the box resulting in a dosage range of 377- 729 mJ cm^-2^ for the test materials. Virus inactivation test results after UV treatment of 8.4 log_10_ PFU of enveloped virus deposited per coupon (8.2 log_10_ PFU for ABS plastic) showed a ≥5 log_10_ PFU inactivation for SS304, polyurethane, and ABS plastic and a ≥3 log_10_ PFU inactivation for NTC and cardboard. As for all other devices, the hardest, smoothest material (SS304) was most effectively treated while the most porous material (cardboard) was hardest to decontaminate.

Overall, the prototype Big Box chamber showed higher virus inactivation compared to almost all other devices, corresponding to the significantly higher UV dosage achieved with the large number of Hg bulbs in the chamber. The data highlights the overall limitations of UV technology to provide complete virus inactivation since virus sterilization was not achieved despite a large, powerful system featuring a total of 320 Philips T8 Hg bulbs.

### Prototype Fixed UV Devices for Room Decontamination

Three prototype devices were also tested that were intended to be installed on the ceiling or wall to provide viral decontamination of the air. These devices followed the same general concept but differed slightly in design and were tested in iterations that featured different UV radiation sources (Hg and KrCl bulbs). Test setup for these devices was largely similar to the previous devices with each test being carried out for five coupons and each inoculated from one of five independent viral preparations. Sterile control and extraction control coupons were also included. However, only one coupon material was tested for each device. Because the purpose of these devices is air decontamination, the specific test material employed here was not particularly important so long as the material was non-porous with high extraction efficiency and the materials provided no additional decontamination properties. SS304 was initially used for testing and was later replaced by quartz glass in one case, as it allows greater UV transmittance to maximize the surface area that would be exposed to UV, and is similar to the way air particles would be exposed at all angles to direct or reflect UV. Testing virus-inoculated glass was a late decision during testing that increased the likelihood of UV penetration from multiple angles in order to prove that UV was able to kill the majority of an 8-log_10_ dried virus challenge. The glass tests also provided time and distance data point that might be used in future iterations of aerosol testing to calculate and predict air flow requirements for air decontamination. Tests were carried out for 5, 10, and 15 s for each device, though the distance from the UV source differed for each device as described below. For all experiments, the devices were powered on for at least 30 min prior to testing to mitigate any start-up fluctuations in UV output.

Devices intended for air decontamination represent a challenge because methods to mimic respiratory enveloped virus for field testing (testing outside of laboratory containment), with a testing turnaround time of 2-weeks , have yet to be developed. While there are nebulization protocols for wet purified virus in laboratory testing, these methods have little practical relevance for field testing of environmentally relevant SARS-CoV-2 virus where the virus is protected by mucus (the surface of which primarily consists of carbohydrate), the infectious particles only consist of ≤0.001% virus, and the infectious 4 um particles are dry, not wet (Hadi et al. 2020; Stadnytskyi et al. 2020). Lab testing of virus suspended in solution demonstrated that virus would not reproducibly survive shipping for field tests (Buhr et al. 2020). For the purposes of this work, the methods for field testing on virus-inoculated surfaces were maintained in order to comparatively screen and assess the effectiveness of the UV bulbs used in the different prototypes, particularly since there was so much variability in dosage and efficacy among different UV sources up to this point. This approach helped with iterative assessments and prototype improvements.

### Prototype Device A (Hg bulb and KrCl bulb iterations)

Prototype device A featured an internal UV source within an enclosed chamber. Fans controlled flow into the chamber where air was exposed to UV-C radiation and then exhausted through vents opposite from the fans. The first prototype contained two Philips TUV 15W/G15 T8 mercury bulbs emitting 254 nm UV-C. Three fans were mounted in the device to provide airflow at 3,030 l min^-1^ total through an effective inner volume of 24.64 l. This led to a residence time of 0.49 s that air would be exposed to UV radiation within the upper chamber. Exposures were at 5 cm, 10 cm, and 15 cm from the UV source.

The second iteration of device A replaced the dual Hg bulb with a single, custom KrCl excimer bulb from Far-UV Sterilay that emitted 222 nm UV-C, with the goal of developing a device with good decontamination efficacy that also posed less of a hazard to personnel exposed to the UV source. The modified prototype A unit was determined to require approximately 5 min for the UV-C output to stabilize. The device was verified to emit a peak wavelength of 222 nm with a slight spike at 252 nm likely from the SiO_2_ glass casing of the bulb (data not shown). The modified device A also included Teflon reflective surfaces to resist dirt build-up and provide reflectance of the UV-C. High purity non-crystalline-fused silica glass plates, also called quartz glass, were added to channel airflow parallel to the UV-C source and increase the total contact time between contaminated air and the UV-C. This increased the total UV dosage applied to air in the unit, thereby providing greater efficacy. The modified prototype device featured three fans providing 1,700 l min^-1^ of airflow each into the unit. One fan was always operated with the UV-C power switch. Two additional power switches were present for each additional fan, therefore, the device could operate at 1,700, 3,400, and 5,100 l min^-1^ airflow. The effective interior volume was 24.33 l.

The efficacy of the UV source within the modified device A (KrCl bulb) was tested with the lid attached. Inoculated quartz glass test coupons were placed individually into a tray and slid inside the unit through slots cut in the frame. Slots were cut at set distances of 4, 10, and 20 cm from the center of the UV bulb. These distances were aligned to prominent design features in the box. The 4 cm test distance (4 cm from the center of the bulb or 2 cm from the edge of the bulb) aligned to an average distance from the bulb in the middle or second air flow channel. The 10 cm distance aligned to the outer channels just behind the quartz glass, and the 20 cm distance also aligned to the outer channels behind the glass at the furthest distance within the device where air would be exposed to UV-C.

### Prototype Device B (Hg bulb and KrCl bulb iterations)

Prototype device B featured a single UV-C source in an open-ended unit. Fans directed airflow into the underside of the unit, and air then exited the frame under and past the UV-C source and then out into the surrounding room air. As with device A, two iterations of the design were tested. The first iteration contained one Philips TUV PL-L 36W/4P Hg bulb. Device B was designed to be mounted to a wall and featured one fan to draw air upwards from underneath the device and exhaust out the top and upper sides. It required a mounting height of 2.15 m in order to ensure that no humans or pets are exposed to the UV-C coming out the sides of the device.

Test exposures for this device were conducted at 5 cm, 10 cm, and 15 cm from the UV source. Coupons were placed in plastic Petri dishes with lids removed for exposures.

The second version of device B contained one KrCl excimer bulb emitting 222 nm UV-C, the same bulb as in the second iteration of device A. With replacement of the 254 nm Hg bulb with 222 nm UV emission, it no longer had the strict requirement of a 2.15 m mounting distance, according to the prototype developer. However, 222 nm UV exposure was still a concern for Navy personnel. Device B contained limited Teflon as a reflective surface was placed near the bulb to direct and concentrate UV outward. Unlike device A, device B does not feature a closed compartment where reflectivity with the Teflon can occur (substantially removing that potential for an increase in applied dosage). The device featured a recessed UV compartment between 10- 15 cm deep with cross sectional area 38.7 cm x 11.4 cm. The compartment was angled upward at approximately 45° from vertical to exhaust air and provide continuous UV exposure of ambient air. The average measured airflow at the compartment outlet was 2,237 l min^-1^. Test exposures for this device were conducted at 5, 15, and 30.5, 61 and 122 cm from the UV source. Coupons were placed in plastic Petri dishes with lids removed for exposures.

### Prototype Device C (Hg bulb type only)

Prototype device C followed a similar concept to device B but with a slightly different configuration and form factor. It was designed to be mounted to a wall and featured one Philips TUV 36W/G36 T8 Hg bulb and two internal fans, with the fans placed to draw air upwards through the unit to exhaust out the top and upper sides. Like device B, it requires a mounting height of 2.15 m in order to ensure that no humans or pets are exposed to the UV-C coming out the upper sides of the device. Test exposures for this device were conducted at 5, 10 and 15 cm from the UV source. Coupons were placed in plastic Petri dishes with lids removed for exposures.

### Mounted Prototypes A, B (Hg bulb and KrCl bulb prototypes), and C (Hg bulb type only) Efficacy

Efficacy results for the mounted prototype A, B and C are shown in **Tables 4** (log_10_ reduction) and **5** (log_10_ survival). Prototypes were tested at representative exposure distances, but the exposure times were much longer than expected for application in order to provide modeling data. Bio-efficacy testing on the original prototypes A, B, and C evaluated the performance of the UV source only. Testing on the modified prototypes A and B evaluated the internal improvements to the device. Thus, bio-efficacy data presented would be significantly less if realistic, shorter times were tested. For the original prototype A with the Hg bulbs, there was minimal log_10_ reduction at different distances and times against virus-inoculated SS304. A modified prototype version with a greatly optimized internal configuration and a KrCl bulb was tested against virus-inoculated quartz glass. Time and cost restrictions prevented a test on virus-inoculated SS304. This modified prototype A unit showed a significant improvement over the original prototype. There were too many significant changes between the first and second prototypes to isolate any single variable as the primary reason for the improved efficacy.

**Table 4.**
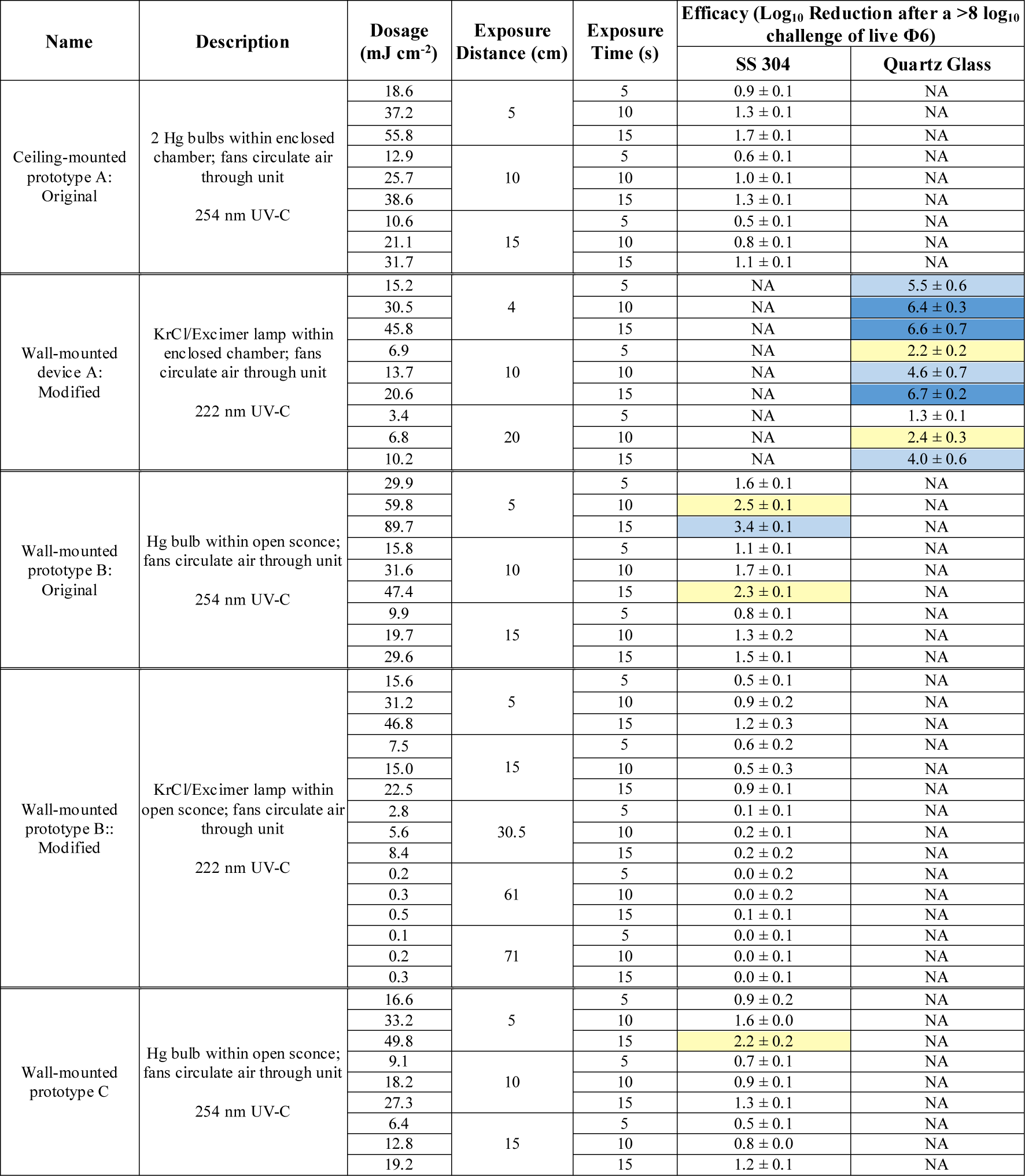
Dosage and efficacy of room-air-irradiating prototypes showing log_10_ reduction data based on the average power from the bulb area, not the peaks. Legend: White (low decontamination) = Fail <2 log_10_; Yellow (sanitation) = Fail ≥2 log_10_, <3 log_10_; Light Blue (disinfection) = Pass ≥ 3 log_10_; and Dark Blue (approaching virus sterilization) = Pass ≥ 6 log_10_. N/A – dosage measurements had no meaning because of the broad-spectrum Xe source.

| Name | Description | Dosage (mJ cm<sup>-2</sup>) | Exposure Distance (cm) | Exposure Time (s) | Efficacy (Log<sub>10</sub> Reduction after a >8 log<sub>10</sub> challenge of live Φ6) |  |
| --- | --- | --- | --- | --- | --- | --- |
|  |  |  |  |  | SS 304 | Quartz Glass |
| Ceiling-mounted prototype A: Original | 2 Hg bulbs within enclosed chamber; fans circulate air through unit<br>254 nm UV-C | 18.6 | 5 | 5 | 0.9 ± 0.1 | NA |
|  |  | 37.2 |  | 10 | 1.3 ± 0.1 | NA |
|  |  | 55.8 |  | 15 | 1.7 ± 0.1 | NA |
|  |  | 12.9 | 10 | 5 | 0.6 ± 0.1 | NA |
|  |  | 25.7 |  | 10 | 1.0 ± 0.1 | NA |
|  |  | 38.6 |  | 15 | 1.3 ± 0.1 | NA |
|  |  | 10.6 | 15 | 5 | 0.5 ± 0.1 | NA |
|  |  | 21.1 |  | 10 | 0.8 ± 0.1 | NA |
|  |  | 31.7 |  | 15 | 1.1 ± 0.1 | NA |
| Wall-mounted device A: Modified | KrCl/Excimer lamp within enclosed chamber; fans circulate air through unit<br>222 nm UV-C | 15.2 | 4 | 5 | NA | 5.5 ± 0.6 |
|  |  | 30.5 |  | 10 | NA | 6.4 ± 0.3 |
|  |  | 45.8 |  | 15 | NA | 6.6 ± 0.7 |
|  |  | 6.9 | 10 | 5 | NA | 2.2 ± 0.2 |
|  |  | 13.7 |  | 10 | NA | 4.6 ± 0.7 |
|  |  | 20.6 |  | 15 | NA | 6.7 ± 0.2 |
|  |  | 3.4 | 20 | 5 | NA | 1.3 ± 0.1 |
|  |  | 6.8 |  | 10 | NA | 2.4 ± 0.3 |
|  |  | 10.2 |  | 15 | NA | 4.0 ± 0.6 |
| Wall-mounted prototype B: Original | Hg bulb within open sconce; fans circulate air through unit<br>254 nm UV-C | 29.9 | 5 | 5 | 1.6 ± 0.1 | NA |
|  |  | 59.8 |  | 10 | 2.5 ± 0.1 | NA |
|  |  | 89.7 |  | 15 | 3.4 ± 0.1 | NA |
|  |  | 15.8 | 10 | 5 | 1.1 ± 0.1 | NA |
|  |  | 31.6 |  | 10 | 1.7 ± 0.1 | NA |
|  |  | 47.4 |  | 15 | 2.3 ± 0.1 | NA |
|  |  | 9.9 | 15 | 5 | 0.8 ± 0.1 | NA |
|  |  | 19.7 |  | 10 | 1.3 ± 0.2 | NA |
|  |  | 29.6 |  | 15 | 1.5 ± 0.1 | NA |
| Wall-mounted prototype B:: Modified | KrCl/Excimer lamp within open sconce; fans circulate air through unit<br>222 nm UV-C | 15.6 | 5 | 5 | 0.5 ± 0.1 | NA |
|  |  | 31.2 |  | 10 | 0.9 ± 0.2 | NA |
|  |  | 46.8 |  | 15 | 1.2 ± 0.3 | NA |
|  |  | 7.5 | 15 | 5 | 0.6 ± 0.2 | NA |
|  |  | 15.0 |  | 10 | 0.5 ± 0.3 | NA |
|  |  | 22.5 |  | 15 | 0.9 ± 0.1 | NA |
|  |  | 2.8 | 30.5 | 5 | 0.1 ± 0.1 | NA |
|  |  | 5.6 |  | 10 | 0.2 ± 0.1 | NA |
|  |  | 8.4 |  | 15 | 0.2 ± 0.2 | NA |
|  |  | 0.2 | 61 | 5 | 0.0 ± 0.2 | NA |
|  |  | 0.3 |  | 10 | 0.0 ± 0.2 | NA |
|  |  | 0.5 |  | 15 | 0.1 ± 0.1 | NA |
|  |  | 0.1 | 71 | 5 | 0.0 ± 0.1 | NA |
|  |  | 0.2 |  | 10 | 0.0 ± 0.1 | NA |
|  |  | 0.3 |  | 15 | 0.0 ± 0.1 | NA |
| Wall-mounted prototype C | Hg bulb within open sconce; fans circulate air through unit<br>254 nm UV-C | 16.6 | 5 | 5 | 0.9 ± 0.2 | NA |
|  |  | 33.2 |  | 10 | 1.6 ± 0.0 | NA |
|  |  | 49.8 |  | 15 | 2.2 ± 0.2 | NA |
|  |  | 9.1 | 10 | 5 | 0.7 ± 0.1 | NA |
|  |  | 18.2 |  | 10 | 0.9 ± 0.1 | NA |
|  |  | 27.3 |  | 15 | 1.3 ± 0.1 | NA |
|  |  | 6.4 | 15 | 5 | 0.5 ± 0.1 | NA |
|  |  | 12.8 |  | 10 | 0.8 ± 0.0 | NA |
|  |  | 19.2 |  | 15 | 1.2 ± 0.1 | NA |

**Table 5.** Dosage and efficacy of room-air-irradiating prototypes showing log_10_ survival data based on the average power from the bulb area, not the peaks. Legend: White (low decontamination) = Fail <2 log_10_; Yellow (sanitation) = Fail ≥2 log_10_, <3 log_10_; Light Blue (disinfection) = Pass ≥ 3 log_10_; and Dark Blue (approaching virus sterilization) = Pass ≥ 6 log_10_. N/A – dosage measurements had no meaning because of the broad-spectrum Xe source.

| Name | Description | Dosage (mJ cm<sup>-2</sup>) | Exposure Distance (cm) | Exposure Time (s) | Log<sub>10</sub> Survival after a >8 log<sub>10</sub> challenge of live Φ6 |  |  |
| --- | --- | --- | --- | --- | --- | --- | --- |
|  |  |  |  |  | Inoculum | SS 304 | Quartz Glass |
| Ceiling-mounted Prototype A: Original | 2 Hg bulbs within enclosed chamber; fans circulate air through unit<br>254 nm UV-C | 18.6 | 5 | 5 | 8.2 +/- 0.0 | 7.3 ± 0.3 | NA |
|  |  | 37.2 |  | 10 |  | 6.9 ± 0.1 | NA |
|  |  | 55.8 |  | 15 |  | 6.5 ± 0.2 | NA |
|  |  | 12.9 | 10 | 5 |  | 7.6 ± 0.2 | NA |
|  |  | 25.7 |  | 10 |  | 7.2 ± 0.3 | NA |
|  |  | 38.6 |  | 15 |  | 7.0 ± 0.1 | NA |
|  |  | 10.6 | 15 | 5 |  | 7.8 ± 0.1 | NA |
|  |  | 21.1 |  | 10 |  | 7.4 ± 0.3 | NA |
|  |  | 31.7 |  | 15 |  | 7.1 ± 0.2 | NA |
| Wall-mounted Prototype A: Modified | KrCl/Excimer lamp within enclosed chamber; fans circulate air through unit<br>222 nm UV-C | 15.2 | 4 | 5 | 8.2 +/- 0.1 | NA | 2.7 ± 1.4 |
|  |  | 30.5 |  | 10 |  | NA | 1.8 ± 0.7 |
|  |  | 45.8 |  | 15 |  | NA | 1.6 ± 1.6 |
|  |  | 6.9 | 10 | 5 |  | NA | 6.0 ± 0.5 |
|  |  | 13.7 |  | 10 |  | NA | 3.5 ± 1.5 |
|  |  | 20.6 |  | 15 |  | NA | 1.4 ± 0.4 |
|  |  | 3.4 | 20 | 5 |  | NA | 6.8 ± 0.3 |
|  |  | 6.8 |  | 10 |  | NA | 5.7 ± 0.7 |
|  |  | 10.2 |  | 15 |  | NA | 4.1 ± 1.3 |
| Wall-mounted Prototype B: Original | Hg bulb within open sconce; fans circulate air through unit<br>254 nm UV-C | 29.9 | 5 | 5 | 8.2 +/- 0.0 | 6.7 ± 0.1 | NA |
|  |  | 59.8 |  | 10 |  | 5.7 ± 0.2 | NA |
|  |  | 89.7 |  | 15 |  | 4.8 ± 0.3 | NA |
|  |  | 15.8 | 10 | 5 |  | 7.1 ± 0.2 | NA |
|  |  | 31.6 |  | 10 |  | 6.5 ± 0.2 | NA |
|  |  | 47.4 |  | 15 |  | 5.9 ± 0.3 | NA |
|  |  | 9.9 | 15 | 5 |  | 7.5 ± 0.2 | NA |
|  |  | 19.7 |  | 10 |  | 6.9 ± 0.4 | NA |
|  |  | 29.6 |  | 15 |  | 6.7 ± 0.2 | NA |
| Wall-mounted Prototype B:: Modified | KrCl/Excimer lamp within open sconce; fans circulate air through unit<br>222 nm UV-C | 15.6 | 5 | 5 | 8.1 +/- 0.3 | 7.6 ± 0.1 | NA |
|  |  | 31.2 |  | 10 |  | 7.2 ± 0.4 | NA |
|  |  | 46.8 |  | 15 |  | 6.9 ± 0.6 | NA |
|  |  | 7.5 | 15 | 5 |  | 7.6 ± 0.2 | NA |
|  |  | 15.0 |  | 10 |  | 7.6 ± 0.6 | NA |
|  |  | 22.5 |  | 15 |  | 7.2 ± 0.1 | NA |
|  |  | 2.8 | 30.5 | 5 |  | 8.1 ± 0.1 | NA |
|  |  | 5.6 |  | 10 |  | 7.9 ± 0.2 | NA |
|  |  | 8.4 |  | 15 |  | 7.9 ± 0.2 | NA |
|  |  | 0.2 | 61 | 5 |  | 8.1 ± 0.2 | NA |
|  |  | 0.3 |  | 10 |  | 8.1 ± 0.2 | NA |
|  |  | 0.5 |  | 15 |  | 8.0 ± 0.1 | NA |
|  |  | 0.1 | 71 | 5 |  | 8.1 ± 0.1 | NA |
|  |  | 0.2 |  | 10 |  | 8.1 ± 0.1 | NA |
|  |  | 0.3 |  | 15 |  | 8.1 ± 0.2 | NA |
| Wall-mounted Prototype C | Hg bulb within open sconce; fans circulate air through unit<br>254 nm UV-C | 16.6 | 5 | 5 | 8.2 +/- 0.0 | 7.4 ± 0.3 | NA |
|  |  | 33.2 |  | 10 |  | 6.7 ± 0.1 | NA |
|  |  | 49.8 |  | 15 |  | 6.1 ± 0.3 | NA |
|  |  | 9.1 | 10 | 5 |  | 7.5 ± 0.2 | NA |
|  |  | 18.2 |  | 10 |  | 7.3 ± 0.2 | NA |
|  |  | 27.3 |  | 15 |  | 6.9 ± 0.2 | NA |
|  |  | 6.4 | 15 | 5 |  | 7.7 ± 0.1 | NA |
|  |  | 12.8 |  | 10 |  | 7.4 ± 0.1 | NA |
|  |  | 19.2 |  | 15 |  | 7.1 ± 0.1 | NA |

For the original prototype B with a Hg bulb, there was minimal log_10_ reduction at different distances and times against virus-inoculated SS304. A modified prototype B with a KrCl bulb emitting 222 nm UV was tested. Test results showed worse efficacy results than the original prototype B. Overall, this device was the least effective of the wall-mounted prototypes, and it was not modified as extensively compared to the modified prototype A. The 222 nm KrCl bulb clearly did not improve efficacy in this prototype.

Prototype C had the least favorable design, and given its low efficacy, it was not pursued for modification.

## DISCUSSION

The focus of this research was to establish reference field test methods for UV decontamination of enveloped virus and to both assess and accelerate improvements in UV devices. Φ6 was selected as a BSL-1, enveloped RNA virus test indicator for both lab and field tests. Φ6 has been widely used as an enveloped virus surrogate (de Carvalgo et al. 2017). It bears structural similarity to many other enveloped viruses including coronaviruses, suggesting that the Φ6 structure should be similarly susceptible to general decontaminants. The structural molecules of the virus are produced by host cells with temperature sensitivity at around 40°C, further suggesting that Φ6 should be similarly susceptible to general decontaminants as animal coronaviruses. The capabilities for measuring UV efficacy using both physics-based equipment and live, enveloped virus test indicators allowed standardized test measurements in both lab and field tests to directly compare the different UV devices.

Greater than 1,000 Φ6 test samples, plus the corresponding Φ6-inoculated, non-treated coupon controls and inoculum controls, were decontamination tested with ≥8 log_10_ Φ6 sample^-1^ during COVID-19 in both lab and field testing (>20 field tests), which included both hot, humid air (Buhr et al. 2020) and this UV testing. A total of 20 independent Φ6 preparations with a titer of 11±0.2 log_10_ were generated on different days by 13 different technicians. Five independent virus preparations were tested on each material for each test. A total of 13 technicians were interchangeably used in different steps of the virus quantitation procedure, and 9 technicians were used on a standard high throughput test day. The reproducibility of the test controls across different test days with numerous technicians and with virus extraction that ranged from 4-14 d after inoculation demonstrated reproducibility of the test method. Methods reproducibility in this data set was critical because coronavirus test methods and results varied significantly during COVID-19, which limited the confidence with which to interpret most published data on coronavirus stability and decontamination (Hadi et al. 2020). Furthermore, critical characterization data needed for accurate and relevant SARS-CoV-2 inactivation testing, such as COVID-19 mucus characterization (Kratochvil et al. 2022), are only now beginning to get published.

Enveloped virus stability had been confirmed previously; purified virus was unstable, but unpurified virus was stable and could be stored dried onto coupons for at least 2 weeks prior to extraction (Buhr et al. 2020). The instability of enveloped virus in solution is a key difference compared to spore quantitation because spores are stable in non-nutrient aqueous solution at temperatures up to at least 65°C. Hence, wet spores can be stored at room temperature for many days alongside inoculated coupons (Buhr et al. 2012), whereas enveloped virus stored at ambient temperature was only stable after drying (Buhr et al. 2020). Furthermore, there was no Φ6 inactivation after unpurified virus was dried onto different surfaces and incubated for a 10 d exposure to 26.7°C at 80% RH, and only 2.4 log_10_ inactivation was seen after treatment at 70°C, 5% RH for 24 h (Buhr et al 2020). More work will be needed to confirm that Φ6 and BSL-2/3 coronaviruses are stabilized similarly in the presence of carbohydrates and mucus, and after drying, but the first challenge is to generate sufficient BSL-2/3 coronavirus to match the titers (and statistical confidence) of the Φ6 tests. This goal has not yet been met. In addition, neither SARS-CoV-2 nor BSL-2 virus field testing is likely to happen with regularity.

The field testing in this manuscript is part of an iterative process of testing, test methods development, and UV analysis. These test methods serve as a baseline screening for military end users, military applications, and for military-relevant environments. To assess manufacturer claims of UV decontamination, different UV bulbs were directly tested with a live/dead enveloped virus assay at >8 log_10_ per test sample with 5 independent virus preps per test material and the virus was dried >24 h prior to testing. Furthermore >50% of inoculated virus was extracted from different control materials for up to 14 d after inoculation and showed to be “live” in a live/dead assay. These controls were needed to validate field test samples. In order to generate confidence and approve a device for end user fielding, additional testing of aerosolized virus will need to be matured. Correlation tests among BSL-1, 2 and 3 enveloped virus will need completed, and then tests will also need to be conducted inside buildings, ships, and aircraft to adjust for variations in air movement and various power outages among all those locations. There will be no lab containment for field testing, so a live/dead assay using a BSL-1 organism will be needed for those tests. As described in the introduction, virus particles will need to be >99.999% non-volatile components, potentially including mucus for a respiratory virus where human airway mucus contains 75-90% carbohydrates (Hadi et al. 2020). Here the assay was simplified using >99.999% sucrose because sucrose is a carbohydrate with representative carbohydrate hydroxyls found on the surface of mucin, a glycoprotein critical to the function of mucus, and because there was not sufficient data to define mucus and drying methods for coronavirus-relevant field testing during COVID-19. Furthermore, sucrose is a known stabilizer that has been historically used for virus purification, and it was empirically determined during Φ6 methods development to positively impact the Φ6 plaque assay (Brakke 1951, 1967; Buhr et al. 2020). Sugars and amino acids also absorb some UV radiation (Uchiho et al. 2015) that partially protects virus in the environment, and this was a secondary benefit to increase confidence in the field test results since mucin is composed of sugars and amino acids.

This UV work was aimed at directly assessing claims that UV radiation would kill enveloped virus using a high confidence live/dead virus assay (≥8 log_10_ of dried virus with virus controls that were recoverable out to at least 2 weeks). Aerosol testing is another layer of testing that will be useful for UV decontamination screening. It is more complex because additional variables such as air movement need controlled and high virus titers are needed in order to generate confidence that the data will represent environmental virus loads. Aerosol testing described in ASTM standard practice E2721-16 was a critical step forward for aerosol testing of wet-dispersed viruses particularly influenza H1N1, and it described one potential artificial saliva recipe that can be added to virus samples (Heimbuch 2013, Anonymous – ASTM 2016). The healthy saliva option contains 0.3% mucin and a total of ∼0.6% non-volatiles, but the ASTM standard practice is flexible and not restricted to that option. This is important because a considerable amount of data describing mucus over the past decade was published after ASTM standard practice E2721-16. Saliva/oral fluids contain mucin glycoprotein, electrolytes, proteins, DNA, lipids, and host cell debris: remnants of dehydrated epithelial and white blood cells (Williams et al. 2006; Lai et al. 2009; Vejerano and Marr 2018; Hadi et al. 2020). Mucin glycoproteins are critical to the function of mucus (Lai et al. 2009), and there are over 20 different types of mucin in humans (Corfield 2015). Similar to sucrose, hydroxyl groups on mucin glycans control water activity (Znamenskaya et al. 2012). Unlike sucrose, the glycan subunits on different mucins are variable, which creates variability in water activity as it pertains to mucin resuspension from dried, commercial sources, and in mucin gel formation (Znamenskaya et al. 2012). Hence, selection of sucrose for field testing during COVID-19 was a simple, homogenous replacement that was easy to standardize for the live/dead screening tests conducted early during COVID-19.

Commercially, two common animal mucins are readily available and frequently used: porcine gastric mucin (PGM) and bovine submaxillary mucin (BSM). PGM is sourced from intestinal epithelium and is desired in experiments due to its sialic acid content that best matches respiratory mucins. BSM is sourced from submaxillary glands in the floor of a cow’s mouth.

While mucin is a major element of saliva, another component of saliva than can interact with mucin is a lung surfactant which is made up of about 90% phospholipids and 10% proteins (Bredberg et al. 2012). The most abundant and active phospholipid in respiratory droplets is dipalmitoylphosphatidylcholine (DPPC) (Niazi et al. 2021). A DPPC saliva recipe (0.9%, sodium chloride, 0.3% porcine gastric mucin, and 0.05% DPPC) was mixed with Φ6, as an enveloped virus surrogate for H1N1, and DPPC protected the mucin to prevent gel formation (Vejerano and Marr 2018). Thus, saliva phospholipids are another potentially important ingredient that might be added to mucus recipes for future iterations of decontamination testing.

The impact of drying on virus stability was a similarly important topic as it is known that enveloped viruses are stabilized by drying (Cox and Wathes, 1995), and SARS-CoV-2 virus has been shown to survive for up to 4 weeks after drying on materials (Riddell et al. 2020). Drying for >24 h was a practical requirement for field testing in this work since samples were routinely shipped to and from test sites. Future aerosol field tests would likely require virus release and monitoring over time, which is beyond the current scope of existing test methods. Defining a final quantity of non-volatiles to generate a standardized mucus solution to mix with enveloped virus is further complicated by the prospect that respiratory disease-afflicted individuals may produce a thicker and/or dryer mucus during different stages of illness as compared to healthy persons (Vejerano and Marr 2018; Kratochvil et al. 2022). Coronavirus particle measurements in 2020 indicated that 12-21 um speech particles would dry to 4 um particles in seconds and smaller particles <1 um in size have such a small probability of containing virus that such a size is likely negligible for infection (Stadnytskyi et al. 2020). A volume/volume calculation indicates that the 12-21 um particles would contain 0.8-3.7% non-volatiles to generate ∼4 um dried particles. Assuming a worst case scenario where the highest level of non-volatiles is protecting enveloped virus, then a mucus recipe with 3.7% non-volatiles contains about 3-6x more non-volatiles compared to existing mucus recipes with 0.6-1.25% total non-volatiles (Aps and Marten 2005; Heimbuch et al. 2013; Anonymous – ASTM 2016; Vejerano and Marr 2018).

COVID-19 mucus characterization was recently published showing a mean load of ∼4.7% solids, and significantly higher amounts of mucin glycoprotein and DNA compared to healthy samples (Kratochvil et al. 2022). The characterization data for COVID-19 mucus non-volatile ingredients was not available when this UV decontamination work was started, but this data will impact future iterations of testing.

A ≥8 log_10_ challenge level was initially set for Φ6 testing because measurements with high concentrations of microbes greatly increase the confidence in inactivation and mitigate the risk of incomplete decontamination (Hamilton et al 2013). Furthermore, coronavirus nasal swabs showed >8 log_10_ virus per swab as calculated using a PCR assay (Leung et al. 2020; Stadnytskyi et al. 2020), and a recent human infection study showed 8.9 log_10_ genomic copies ml^-1^ from nasal samples 5 d after infection (Killingley et al. 2022). The limitation of genomic sampling is that it does not equate to infectious particles. However, an individual highly infected with SARS-CoV-2 can emit >8 log_10_ virus particles in a 24 h period (Stadnytskyi et al. 2020). Extrapolation of that data indicates that multiple personnel within confined spaces, limited ventilation, and low humidity may generate considerably >8 log_10_ of infectious virus at any given point in time, and is a critical considerations in both military and transportation applications. The infectious particle data further justifies the need for ≥8 log_10_ challenge regardless of whether the virus is aerosolized or on surfaces.

UV radiation does not fall under the United States Environmental Protection Agency (EPA) jurisdiction for disinfection claims since it is not classified as a chemical disinfectant. Nonetheless, for this study that began in 2020, the inactivation goal was reduced from ≥7 log_10_ to ≥3 log_10_ inactivation during the COVID-19 pandemic to match the EPA N-list for decontaminants. This was initially helpful because inactivation numbers for sanitation, disinfection, and sterilization (Rutala et al. 1996) could be used for assessments of decontaminants including UV. However, the 3 log_10_ kill threshold is now in question because a very recent human infection study showed that only 10 TCID_50_ of wild-type SARS-CoV-2 (equivalent to ∼7 PFU) was needed to infect >50% of unvaccinated humans in a 36-volunteer group with no prior infection (Killingley et al. 2022). This human infection study validates the original quantitative objective to show enveloped virus inactivation of ≥7 log_10_ out of a ≥8 log_10_ challenge in order to increase decontamination confidence. These high kill objectives/requirements are important to validate decontamination claims for respiratory pathogens, which spread rapidly and continuously through air and likely also through surface transfer. The wide variability in virus preparation and test methods and the associated results published during COVID-19 has decreased end user confidence levels for evaluating decontaminants (Hadi et al. 2020). The limitation of direct SARS-CoV-2 virus testing is the lack of statistically significant test data with multiple independent preparations of virus at high virus titers (>10 log_10_ of virus ml^-1^ of culture medium at the time of virus harvest and without virus concentration or purification). These quantitative requirements need addressed in order to generate end user confidence that data can be reproduced (statistical accuracy) and that virus titers can match the virus stabilization and quantities that may be encountered in the environment.

This background data indicates that detailed methods for mucus and aerosol testing of respiratory coronavirus virus will need to be defined, characterized, and developed for future iterations of decontamination testing. These should also include correlation testing among Biosafety Level 1, 2, and 3 enveloped viruses. Data from methods with increasingly higher confidence will improve the likelihood of end user acceptance of UV technologies. The work here showed that enveloped virus could be field tested at >8 log_10_ of stabilized enveloped virus using multiple preparations of virus and a live/dead assay. The quantitation could be reproduced and field tested with a 2-week data turnaround to provide an initial UV screening/selection. The UV test results here also showed that high UV dosages are needed to inactivate enveloped virus protected by environmental debris, and porous materials are difficult to decontaminate, particularly in comparison to purified virus alone. These limitations of UV are well documented by regulatory agencies, and those limitations also apply to SARS-CoV-2 (United States Food and Drug Administration 2021; United States Environmental Protection Agency 2021). Nonetheless, UV efficacy was measurable and very high dosages were effective even on relatively porous materials like cardboard. It is unlikely that UV would be useful for highly porous fabrics used to make bags, carpeting, and clothing, and those were not tested. In contrast, hot, humid air inactivates debris-laden microbes with similar kinetics regardless of material porosity (e.g. Buhr et al 2012, 2015, 2016, 2020). This is a hallmark difference between highly penetrative decontaminants and a surface decontaminant like UV.

Anti-viral efficacy among the different UV devices ranged from no decontamination up to nearly achieving enveloped virus sterilization. Enormous variability in dosage and efficacy was measured within and among the different devices. The prototype medium conveyer generated the highest virus inactivation per dosage. Inoculated coupons were exposed to UV-C on three sides since the coupons were set on a flat surface during exposure in the conveyer. The big UV box also generated high levels of virus inactivation, but the medium conveyer was highest efficacy dose^-1^. In contrast the handheld devices, pulsed Xenon devices, LEM, and the original prototypes A, B, and C, and modified prototype B were all evaluated with a UV source emitted from predominantly one direction with slightly varying angles of exposure. The increased angles of exposure in the conveyer and big box likely improved UV-C penetration.

Hence, the unique geometry, design, and electronics of each device impacted the effectiveness above and beyond the wavelength and dosage.

The variability seen among the various tested devices strongly indicates that all UV devices need to be measured for both UV dosage and for anti-viral efficacy before they are incorporated into decontamination protocols. The efficacy of a pulsed Xe bulb was measurable at close distances, but significantly lower than Hg bulbs. Pulsed Xe devices do have some practical advantages such as requiring minimal warm up time and no Hg toxicity. LEDs have the lowest hazard and lowest variability in UV output. However, the availability of UV LEDs has been limited, and UV dosages can also be limiting depending on the manufacturer, model, and the electronics and overall design of any given device. Longer wavelength UV (272 nm) showed the best efficacy in handheld devices, and 272 nm is more penetrating than short wavelengths.

The 222 nm KrCl sources showed measurable efficacy in conjunction with proprietary prototype advancements. Additional testing with >8 log_10_ debris-laden virus is needed because the 222 nm KrCl testing was limited due to time and cost constraints.

Finally, decontamination with UV comes with tradeoffs that affect the decision of the end user. The time of exposure needed to generate efficacy needs to be assessed by end users because long exposure times will limit the utility of UV, especially for handheld and air decontamination devices. Use of handheld UV devices is also very hard to standardize, which increases safety risks for end users (United States Food and Drug Administration 2021b). Another tradeoff to be assessed by end users is the need for cleaning and maintenance of UV devices to remove dirt and debris that accumulates on the radiation sources, and/or to change radiation sources. Devices and methods to monitor UV dosage over time are needed to assist in maintenance, a particularly important subject that is rarely addressed. Additional tradeoffs are ozone generation, which reached toxic levels up to 3.6 times higher than OSHA limits for some devices (Claus 2021), and operation times; Hg bulbs, in particular, require warmup times in order to reach a steady-state. In general, Hg bulbs generate a maximum intensity quickly, and then the intensities were stabilized at a lower level after a warm up period. The Hg devices would have performed better had only this initial dose been tested, but that data would not translate to practical application because it will be hard to standardize turn-on times for end users, especially for handhelds. Lastly, the end user needs an understanding of the organism(s) to be killed, how it is stabilized in the environment, and the impact of test methods on results, as these factors will impact confidence in any application. Assessment of these tradeoffs will facilitate practical application of UV decontamination. UV has potential for augmenting current practices for limiting the spread of enveloped virus, but UV cannot and should not replace normal cleaning and hygienic practices or air filtration and ventilation. Equipment breakdowns/failures, electrical outages, and maintenance shortcomings are common, normal, and realistic. Hence, coronavirus test methods need to significantly improve in order to approximate realistic environmental conditions with debris-laden, dried, high titer virus and increase UV test confidence. As test standards, UV field validation methods, and UV sources improve, UV might become a more viable option for augmenting decontamination in some applications. Multiple layers of testing, including field validation testing such as that described herein, will be needed because of the extraordinary variability of UV output from different devices. In addition, methods to continuously monitor UV output and maintenance and cleaning guidelines are critical areas that need addressed.

## ACKNOWLEDGEMENTS

This work was supported through funding and program support provided by Naval Sea Systems Command, Defense Innovation Unit, Naval Advanced Medical Devices, and the Defense Threat Reduction Agency (DTRA), Hazard Mitigation Capability Area (BA2 and 3 funds, Project Number CB10141). We thank Rich Wiersteiner, Jon Cofield, Heather Ichord, Janet Weir, Glenn Lawson, Chuck Bass, James Noah, John Aaron Miller, and Joe Schumer for support. We thank Jason A. Fallen, Dr. Joseph Hunt, Carlos Murillo, Kira Baugh, and Julie Caruana for outstanding technical support and/or assistance with editing the paper. This manuscript was approved for public release on 1/27/2022 as #6156; NSWCDD PN-22-00021 and 8/1/2022 as #6232; NSWCDD PN-22-00131 and will be released as a pre-print at bioRxiv (Buhr et al.).

